# Distinct Transcriptomic Biomarkers and Pathways Associated with Cardiovascular and Neurovascular Dysregulation in Long COVID-19 Brain Fog

**DOI:** 10.1101/2025.06.10.656785

**Authors:** Orusa Ashfaque, Harsh Kumar, Mohit Mazumder

## Abstract

**Background:** Long COVID-19 often manifests as persistent cognitive impairments, commonly referred to as brain fog, with poorly understood biological mechanisms. Previous studies highlighted neuroinflammation and blood-brain barrier disruption. Here, we identify distinct biomarkers and dysregulated pathways, with a focus on cardiovascular and neurovascular involvement.

**Methods:** We analyzed public RNA-sequencing data (GEO: GSE251849) comparing long COVID-19 patients with brain fog, long COVID-19 patients without cognitive symptoms, convalescent patients, and healthy controls. Differential gene expression, principal component analysis, and pathway enrichment analyses were performed using DESeq2, KEGG, GO, and Enrichr tools.

**Results:** Our analysis revealed a distinct gene expression profile in patients with brain fog, identifying exclusive biomarkers such as *NRCAM, GRIN2C, NOS2,* and *EDNRB.* Pathway analysis revealed significant dysregulation in cardiovascular, calcium signaling, and relaxin signaling pathways, suggesting novel biological mechanisms that contribute to cognitive impairment. These findings expand beyond neuroinflammation and highlight potential cardiovascular involvement in long COVID-related cognitive dysfunction.

**Conclusion:** Our study reveals novel transcriptomic signatures that highlight cardiovascular and neurovascular dysregulation, providing promising diagnostic biomarkers and therapeutic targets. Future research should validate these findings and investigate therapeutic interventions addressing these unique pathways.

## 1. Introduction

COVID-19, caused by severe acute respiratory syndrome corona virus 2 (SARS-CoV-2), emerged in China in December 2019 and rapidly spread worldwide, leading to a global pandemic resulting in millions of deaths (Rodriguez-Morales *et al*., 2020). It is highly infectious and spreads through respiratory droplets produced by coughing and sneezing. Common symptoms include fever, cough, fatigue, and shortness of breath, with an incubation period of 2–14 days (Kishor and Ramhari, 2020). The spike protein of the virus binds to angiotensin-converting enzyme 2 (ACE2) receptors with high affinity, facilitating viral entry and causing respiratory symptoms. In severe cases, cytokine activation leads to sepsis and acute respiratory distress syndrome (Hon *et al*., 2020; Rao and Bhattamisra, 2020). Diagnosis relies primarily on RT□PCR testing of respiratory samples, with imaging techniques such as chest X-rays supporting diagnosis in atypical cases (Vishal *et al*., 2024). Treatment strategies include supportive care, vaccination, antiviral therapy (e.g., remdesivir and poxolovid), and immunomodulators (e.g., tocilizumab) for severe cases. Monoclonal antibodies have been used for passive immunization (Vishal *et al*., 2024).

The complications and consequences of COVID-19, including long COVID-19, are diverse and impactful. Long COVID-19, also known as postacute sequelae of SARS-CoV-2 infection, affects a significant proportion of patients, with symptoms persisting for weeks or months after the acute phase (Golzardi *et al*., 2024; Bellanti, 2023). These symptoms involve various organ systems, including the neuropsychiatric, cardiovascular, respiratory, immune, renal, endocrine, gastrointestinal, and dermatological systems (Golzardi *et al*., 2024). Multiple cases of long-term complications associated with post-COVID-19 infection have emerged, including arterial thrombosis, venous thromboembolism (VTE), cardiac thrombosis, lung fibrosis, brain fog, stroke, mood dysfunction, and dermatological complications (Desai *et al*., 2021; Lee *et al*., 2021). Long COVID-19 symptoms include fatigue; brain fog symptoms such as sluggish thinking; memory issues; difficulty concentrating; respiratory issues such as shortness of breath; cough; neurological symptoms such as headache, dizziness, and loss of smell or taste; gastrointestinal and cardiovascular symptoms such as diarrhea, chest pain, and pulmonary embolism; and other symptoms such as rash, arthralgia, depression, and anxiety (Van Der Feltz-Cornelis *et al*., 2024; Mathew *et al*., 2021; Bellanti, 2023).

Among these, cognitive dysfunction, often termed “brain fog”, has been widely reported. Epidemiological studies have estimated that long-term cognitive impairment occurs in approximately 17–28% of COVID-19 survivors (Möller *et al*., 2023). The brain fog describes cognitive dysfunction, including concentration problems, word-finding difficulties, poor memory, and disorientation. Although it is not a recognized medical diagnosis, it can be a debilitating manifestation of various conditions, such as stress, diet, sleep deprivation, and physical or mental illness. It can occur in many medical contexts but has been especially noted in association with long COVID-19, often leading to impaired functioning or distress (Möller *et al*., 2023; Krishnan *et al*., 2022).

Understanding the prevalence of brain fog is crucial, as the epidemiology of COVID-19 indicates a substantial burden among individuals recovering from SARS-CoV-2 infection. A meta-analysis revealed an overall pooled prevalence of approximately 43%, including 54% of hospitalized patients and 34% of non-hospitalized individuals who experienced long-term symptoms. Regional differences exist, with prevalence rates of 51% in Asia, 44% in Europe, and 31% in North America (Su *et al*., 2023). Global estimates suggest that approximately 6.2% of symptomatic patients exhibit at least one long COVID-19 symptom cluster at various time points postinfection. In some cohorts, even greater proportions of patients reported lingering memory, attention, and executive function deficits months after acute infection (Grover *et al*., 2021; Ceban *et al*., 2021). These findings underscore the urgent need to explore the biological mechanisms underlying brain fog.

The clinical presentations of brain fog include memory lapses, difficulty concentrating, slowed information processing, and general mental fatigue (Krishnan *et al*., 2022). Emerging evidence suggests that these symptoms are associated with neuroinflammation and cerebrovascular dysfunction (Talkington *et al*., 2025). Autopsy and biomarker studies have revealed signs of central nervous system (CNS) involvement in long-term COVID-19 patients with cognitive complaints. For example, markers of astrocyte injury and blood–brain barrier (BBB) disruption, such as glial fibrillary acidic protein (GFAP) and S100β, are elevated in patients experiencing brain fog (Talkington *et al*., 2025). Similarly, a study by Greene *et al*. (2024) reported that long-term COVID-19 patients with cognitive impairment exhibit persistent BBB breakdown and sustained systemic inflammation. Neuroimaging and histopathology suggest that even mild SARS-CoV-2 infection can trigger microglial activation and white matter changes reminiscent of “chemo-brain”, leading to cognitive deficits weeks to months later (Fernández-Castañeda *et al*., 2022).

Although systemic inflammation and BBB leakage have been implicated, the specific molecular pathways and gene networks underlying long-term COVID-19-related cognitive symptoms remain unclear. Here, transcriptomic profiling provides a promising approach. Peripheral blood, which is easily accessible, may reflect ongoing immune and inflammatory processes affecting the brain (Missailidis *et al*., 2024). High-throughput RNA sequencing enables unbiased identification of differentially expressed genes (DEGs) in blood cells that could serve as molecular fingerprints of long COVID-19 brain fog (Missailidis *et al*., 2024).

Prior work has demonstrated that post-COVID-19 patients can exhibit distinct immune gene expression profiles. For example, one transcriptomic study revealed that long COVID-19 patients with brain fog presented upregulation of T-cell activation and angiogenesis genes and downregulation of platelet and coagulation genes relative to recovered individuals (Greene *et al*., 2024). Such gene expression alterations can point to dysregulated pathways, such as TGF-β signalling and HIF-1α signalling for hypoxia, that might contribute to cognitive deficits (Greene *et al*., 2024). Molecular signatures associated explicitly with cognitive symptoms can be isolated by comparing gene expression between long COVID-19 patients with and without brain fog and among convalescent patients (Kayalar *et al*., 2024).

Thus, among the myriad of long-term COVID-19 symptoms, this study focuses on elucidating the molecular basis of brain fog, a prevalent yet poorly understood condition, by identifying distinct transcriptomic signatures.

The objective of the present study was to leverage a publicly available transcriptomic dataset (GEO: GSE251849) of peripheral blood samples from long COVID-19 patients with and without brain fog, convalescent COVID-19 patients, and healthy controls (Greene *et al*., 2024). We performed differential gene expression analysis via DESeq2, along with Kyoto Encyclopedia of Genes and Genomes (KEGG) and Gene Ontology (GO) enrichment analyses, to interpret the functional significance of the DEGs. Principal component analysis (PCA) was employed to visualize gene expression patterns across cohorts, and a Venn diagram comparison of DEGs helped identify genes uniquely dysregulated in the brain fog group. Our primary aim was to identify candidate biomarkers and pathways that might explain cognitive symptoms in long-term COVID-19 patients and could serve as diagnostic indicators or therapeutic targets. COVID-19’s long-term cognitive consequences, commonly termed brain fog, impact a significant portion of recovering patients. Recent studies identified neuroinflammation and compromised blood-brain barrier integrity as underlying mechanisms. However, there remains limited exploration into cardiovascular and neurovascular involvement, which could significantly contribute to these cognitive symptoms. This study aims to identify unique transcriptomic biomarkers and pathways specifically related to cardiovascular and neurovascular dysregulation in long COVID-19 brain fog.

By identifying these unique transcriptomic signatures, our study provides insight into the biological correlates of brain fog in long-term COVID-19 patients. Such knowledge is critical for developing objective diagnostics and guiding future interventions to alleviate these disabling neurological symptoms.

## 2. Methodology

### 2.1 Data extraction

We collected datasets from the Gene Expression Omnibus (GEO) of the National Center for Biotechnology Information (NCBI). Public RNA-seq data (GSE251849) comprising peripheral blood mononuclear cells (PBMCs) from four groups (brain fog, long COVID without cognitive issues, convalescent, healthy controls) were analyzed. This dataset includes RNA sequencing data from peripheral blood mononuclear cells (PBMCs) derived from individuals classified into four groups: healthy controls, convalescent individuals (recovered from COVID-19 patients), long COVID-19 patients without brain fog, and 12-week persistently symptomatic long COVID-19 patients. This dataset was particularly appropriate for our study objectives because it allowed direct comparisons between patients with and without brain fog, thereby enabling the identification of transcriptomic signatures uniquely associated with cognitive dysfunction in long COVID-19 patients.

### 2.2 Data Cleaning & Preparation

The raw gene expression data were downloaded as a compressed file, GSE251849_Counts.txt.gz, available in the supplementary section of the GEO page. This file contains read counts for 62,710 genes across all the samples. The raw count matrix was imported into Microsoft Excel and transposed to align genes as rows and samples as columns for downstream analysis. Sample metadata (disease group) were appended to each column to facilitate grouping and labelling during further analysis. To ensure data quality and prepare the dataset for differential gene expression and dimensionality reduction, a multistep cleaning and normalization process was conducted:

**1. Normalization:** Log2 transformation was applied to each count value (after adding a pseudo count of 1) to bring the data to a comparable scale for visualization and PCA. Any nonnumeric or undefined entries (e.g., “#NUM!”) were replaced with zeroes.
**2. Filtering Low-Quality Data:** Genes with total expression counts less than 10 across all 23 samples were excluded to remove background noise and low-expression artifacts. This filtering reduced the dataset from 62,710 genes to **23,731 relevant genes**.
**3. Duplicate and missing values:** Duplicate gene entries and missing values were identified and removed. The final cleaned dataset consisted of only uniquely mapped, biologically meaningful genes with adequate expression.

### 2.3 Dimensionality reduction

Principal component analysis (PCA) was conducted via the T-Bioinfo server (https://server.t-bio.info/) to investigate the data patterns. The unsupervised analysis pipeline was executed via the pipeline to generate the PCA plot, as illustrated in **Figure 1**. Initially, normalized data comprising four groups were uploaded, with the respective data allocated to each group. Groups A, B, C, and D consisted of samples from healthy individuals, convalescents, and long COVID-19 brain fog, respectively. The files were subsequently submitted. To commence the unsupervised analysis, the process was initiated by selecting ‘Start’, followed by the PCA_R Library, where the number of principal components was set to three, and the data were scaled and transposed. The pipeline was designated PCA_Long_COVID, and the analysis was executed on the cluster.

**Figure 1:**
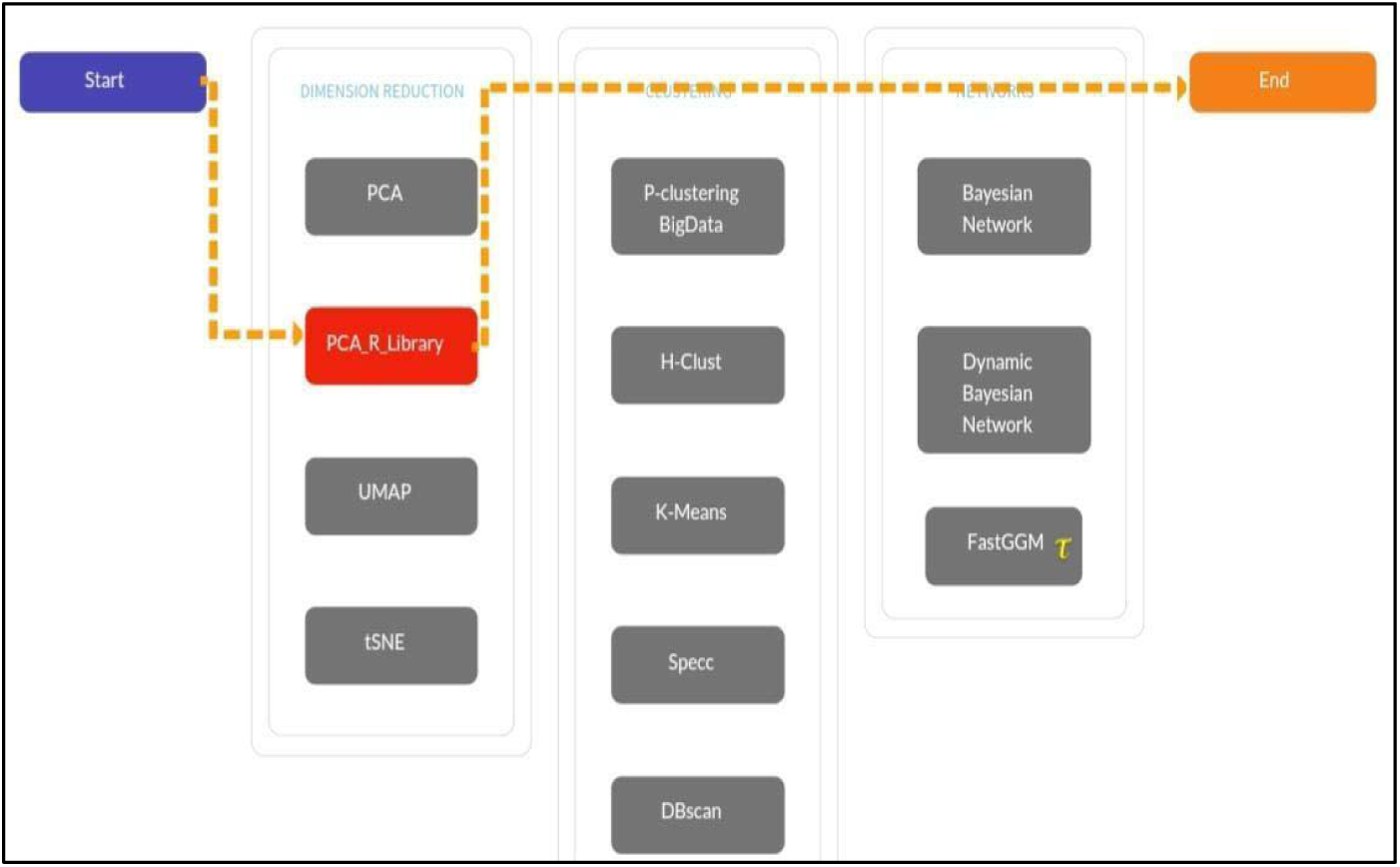
Workflow for the principal component analysis (PCA) pipeline using the T Bio Info server. This figure outlines the steps of the unsupervised analysis pipeline followed to generate PCA plots for differentiating between healthy, convalescent, long COVID-19, and long COVID-19 and brain fog groups.

### 2.4 Analysis of differentially expressed genes (DEGs)

Gene expression analysis was conducted via the T-bioinfo server to examine differential gene expression. **Figure 2** illustrates the pipeline employed for this analysis. Initially, a table containing raw read counts of gene expression data was uploaded. The dataset labelled “Raw Long_Covid” was first uploaded, followed by the selection of two groups: Differential Groups A and B, comprising samples from control and long COVID-19 patients. A differential expression graph was generated, and statistical analysis was performed via DESeq2. The parameters included a count filter for DESeq2 at 9, a volcano plot log fold change (LogFC) threshold of 2, a volcano plot p value threshold of 0.05, and the database was specified as human. Similar procedures were applied to analyse the following comparisons: 1) control vs. convalescent, 2) control vs. long COVID-19 without brain fog, and 3) control vs. long COVID-19 with brain fog. A text file containing unfiltered raw data from DESeq2 was opened in an Excel file, and filters were applied for values greater than 2 and less than −2, with a p value and adjusted p value (padj) less than 0.05. These steps were repeated for all groups, including long COVID-19 only, brain fog only, and convalescent against healthy controls, with minor adjustments to the filter parameters.

**Figure 2:**
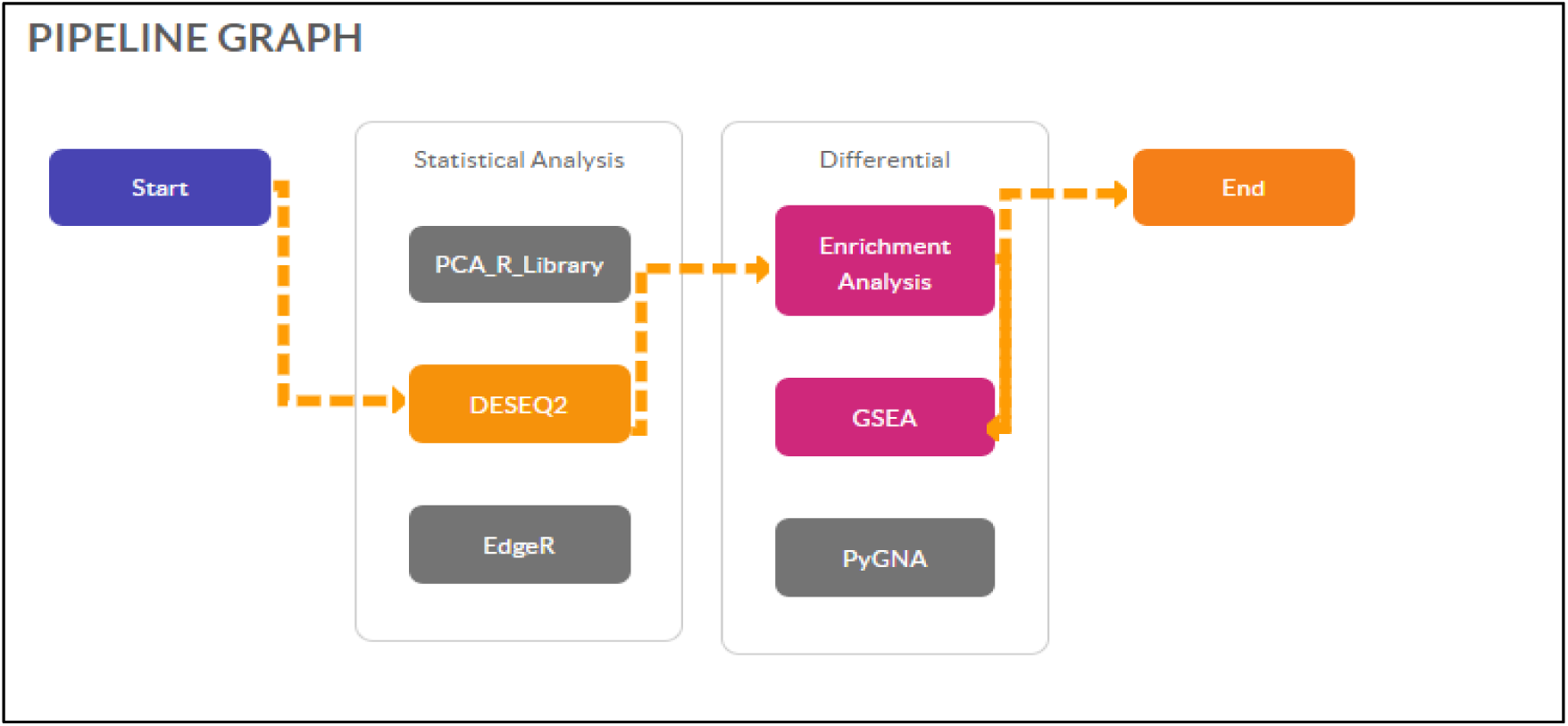
Workflow for the differential gene expression (DGE) analysis pipeline. The figure illustrates the analytical pipeline used to analyse differential expression between the control and various COVID-19-related conditions via the T-Bio Info platform.

### 2.5 Differential gene enrichment analysis

Subsequently, differential gene enrichment analysis was conducted to identify biological pathways and functional categories associated with the differentially expressed genes (DEGs), as depicted in **Figure 2**. Gene Ontology (GO) and Kyoto Encyclopedia of Genes and Genomes (KEGG) pathway enrichment analyses were performed on the statistically significant DEGs. An adjusted p value threshold of 0.05 was applied to determine enrichment significance, ensuring the retention of only biologically relevant terms. The false discovery rate (FDR) method was employed to correct for multiple hypothesis testing. Before enrichment analysis, DEGs were filtered on the basis of an adjusted p value of less than 0.05 from the DESeq2 output, ensuring the inclusion of only robust and statistically reliable DEGs in the GO and KEGG analyses.

### 2.6 Comparative analysis of DEGs and pathways

Comparative analysis of DEGs and pathways was facilitated via Venn diagrams to visualize the overlap between multiple gene lists. The Venny tool was utilized to input the genes, observe the Venn diagram, and download the list of exclusive and common genes among healthy controls (H), convalescent individuals (Con), long COVID-19 patients without brain fog (LC), and long COVID-19 patients with persistent cognitive symptoms (brain fog) (BLC). This approach aids in identifying shared and exclusive genes across different experimental conditions, time points, or differentially expressed gene sets.

### 2.7 Pathway and GO analysis of exclusive DEGs via Enrichr

Pathway and GO analyses of exclusive DEGs from long COVID-19 patients with brain fog (LCBF) were conducted via Enrichr, a web-based tool for gene set enrichment analysis. Enriched KEGG pathways and Gene Ontology (GO) terms were identified on the basis of a significance threshold of p<0.05. Clustergrams were generated to visualize the associations between input genes and enriched terms, where rows represent input genes, columns represent the top enriched terms (ranked by enrichment score), and cell highlights indicate gene-term overlaps.

### 2.8 Identification of novel biomarkers

Our study aimed to identify potential exclusive biomarkers with diagnostic or prognostic value in LCBF by examining differentially expressed genes (DEGs) and enriched pathways. The upregulated, downregulated, and exclusive DEGs were cross-referenced with a literature review to assess the novelty of specific genes in the context of LCBF.

## 3. Results

### 3.1. Data Extraction

A pivot table was created from a series matrix file, which helped summarize the dataset and identify the number of samples and the types of groups involved in the study. **Table 1** summarizes the entire dataset via a pivot table. A total of 23 samples were included in the study, with participants distributed across four groups.

**Table 1:** Pivot table, distribution of 23 participants across four clinical groups: Healthy(n=7), COVID-19 Convalescent (n=5), Long COVID-19 without brain fog (n=6), and Long COVID-19 with brain fog (n=5). The sex distribution is shown, with 19 females and four male participants.

| Count of Samples | Column Labels |  |  |
| --- | --- | --- | --- |
| Row Labels | Female | Male | Grand Total |
| COVID-19 convalescent | 3 | 2 | 5 |
| Healthy | 6 | 1 | 7 |
| Long COVID-19 | 5 | 1 | 6 |
| Long COVID-19 brain fog | 5 |  | 5 |
| **Grand Total** | **19** | **4** | **23** |

### 3.2 Dimensionality reduction

Principal component analysis (PCA) **(Figure 3)** revealed unique gene expression patterns in long COVID-19 brain fog, forming a distinct cluster from the other groups (PC1: 20.78%, PC2: 9.38%, PC3: 8.45%). Healthy controls were separated from the disease groups, with Convalescent and Long COVID-19 patients showing some shared characteristics.

**Figure 3:**
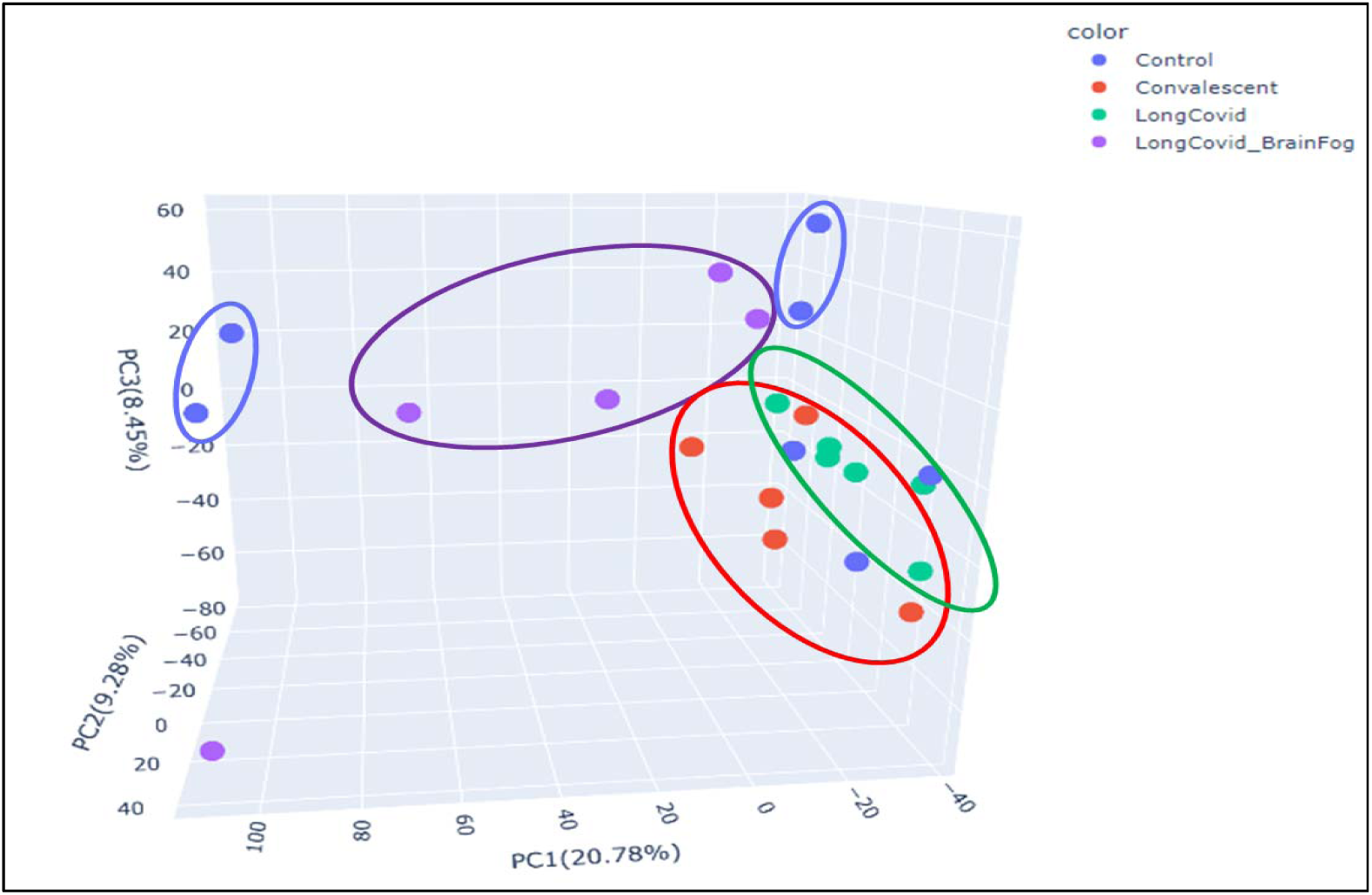
The x-axis represents the first principal component (PC1), accounting for 20.78% of the variance. The y-axis represents the second principal component (PC2), accounting for 9.38% of the variance. The z-axis represents the third principal component (PC3), accounting for 8.45% of the variance. The long COVID-19 brain fog group is a distinctly separated cluster, indicating unique features or biomarkers.

### 3.3 Differential gene expression analysis and functional characterization

#### 3.3.1 Healthy vs long COVID-19 only (LC only)

##### 3.3.1.1 Top upregulated and downregulated DEGs

Differential gene expression analysis revealed several genes whose expression significantly changed. The total number of DEGs in the long COVID-19-only group was 262, among which 66 were upregulated genes and 196 were downregulated genes. The *IGHV7-4-1* gene was notably upregulated among the long COVID-19 samples. Among the downregulated genes, *CHMP1B-AS1* presented the greatest negative fold change (–25.09), while *GJB2* and *OLR1* were also significantly downregulated. **Table 2** presents the top upregulated and downregulated genes, highlighting the distinct molecular profiles associated with long COVID-19.

**Table 2:**
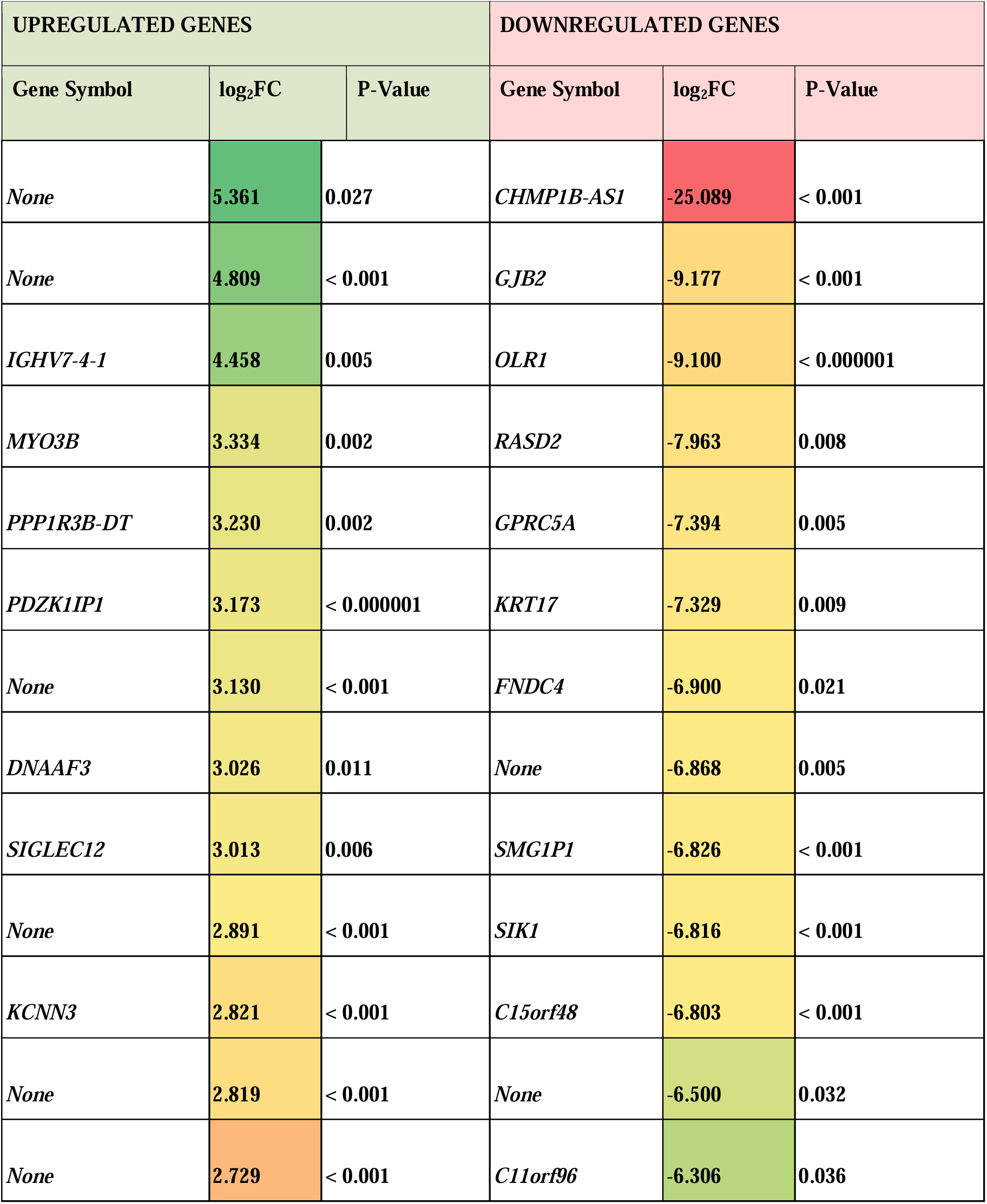
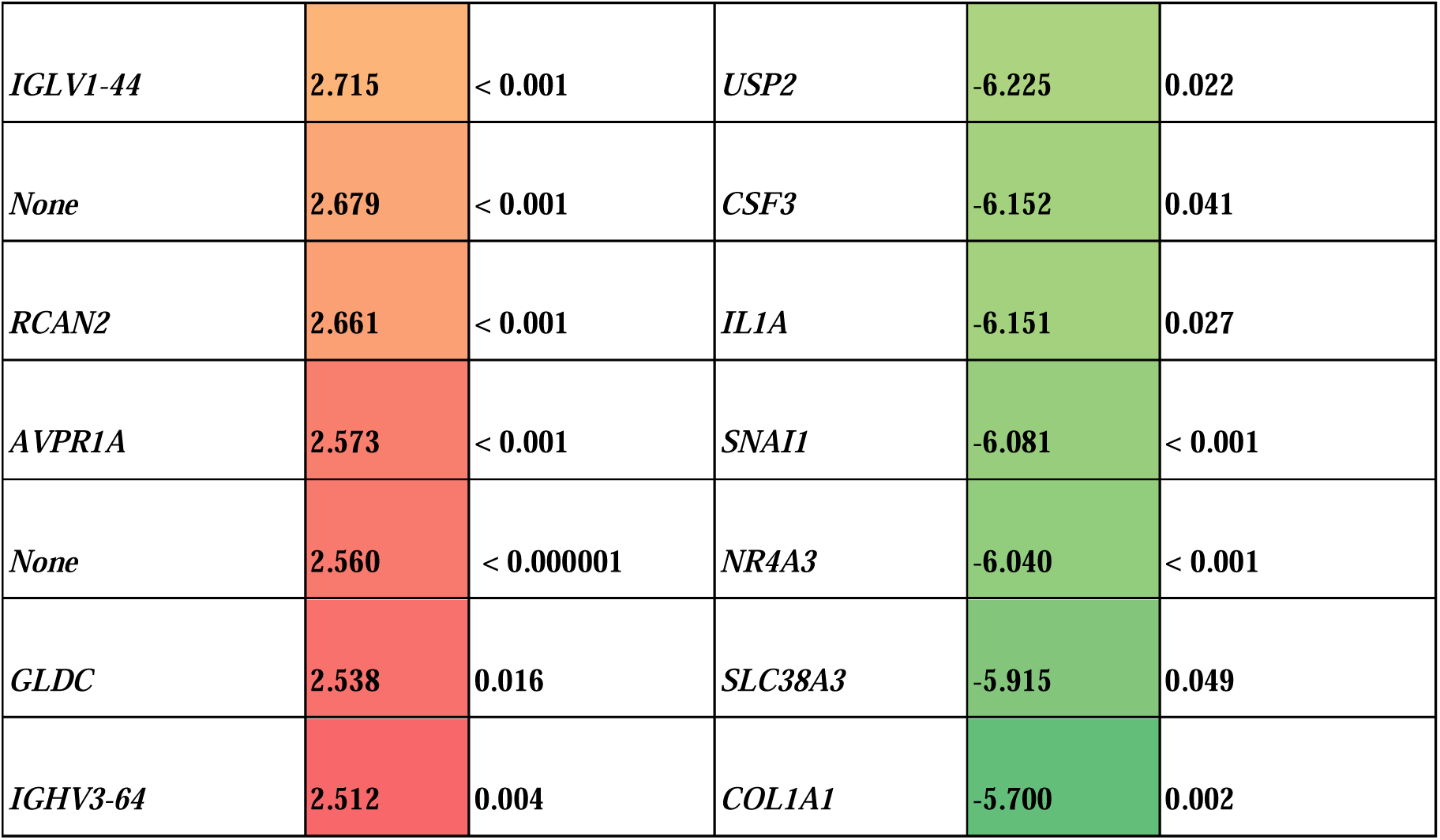
Top upregulated and downregulated DEGs in the Long COVID-19-only (LC) group.

##### 3.3.1.2 Volcano plot and heatmap for the long COVID-19-only (LC-only) group

The volcano plot (**Figure 4A)** indicates significant changes in gene expression, with *PDZK1IP1* being notably upregulated, whereas *SMG1P1* and *OLR1* are the genes whose expression is most downregulated. The heatmap (**Figure 4B)** shows that *OLR1* and *FOXJ1* were overexpressed in the control samples, whereas in all Long COVID-19 samples, *OLR1, FOXJ1*, and *SMG1P1* were consistently downregulated, with the highest number of differentially expressed upregulated genes observed in the LongCOVID007 sample.

**Figure 4:**
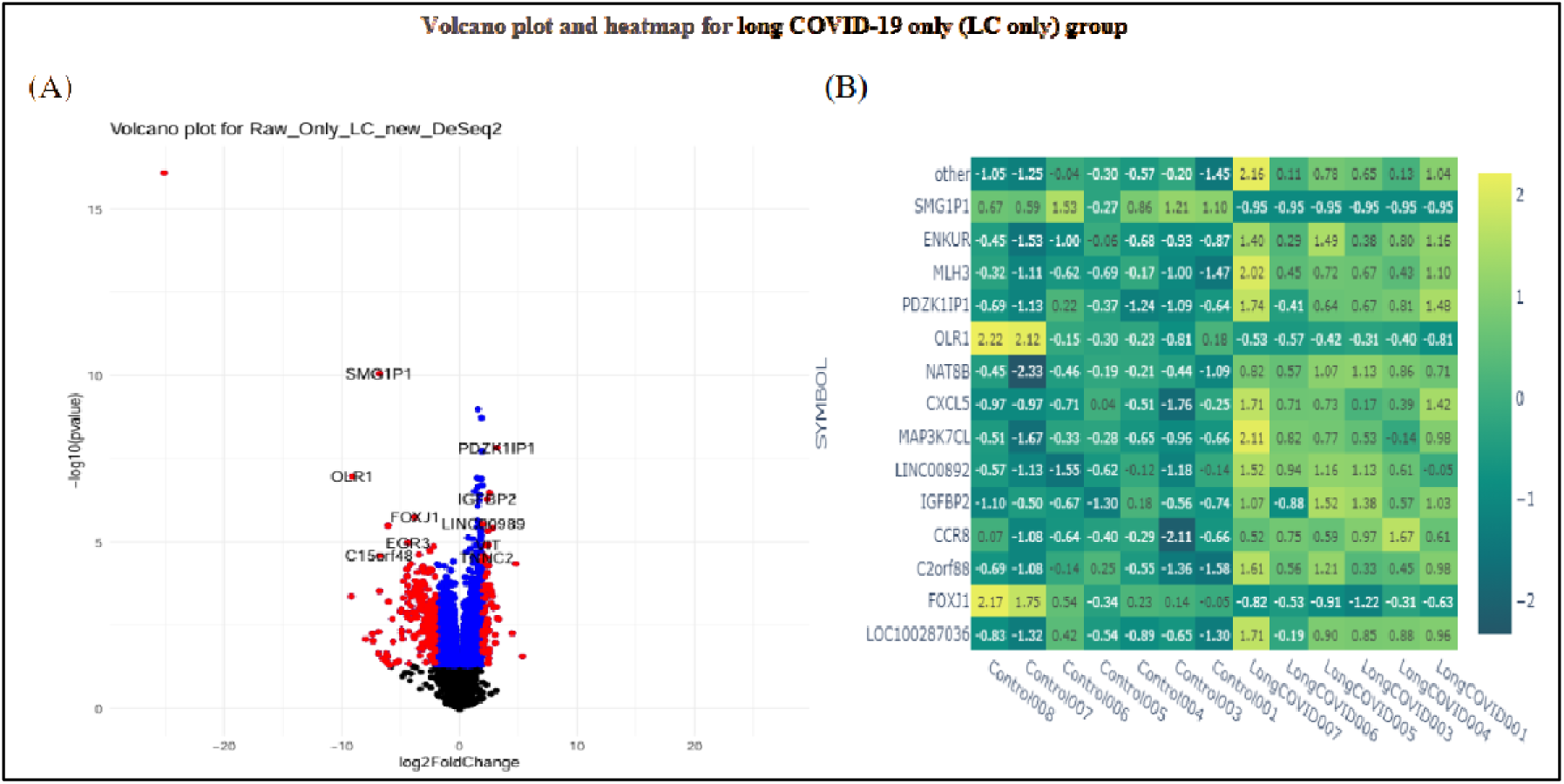
Differential gene expression analysis across the control and Long COVID-19 (LC)-only groups. **A.** Volcano plot showing the differential gene expression between the control and long COVID-19 samples. The upregulated genes are displayed on the right, and the downregulated genes are on the left, with significance indicated by the -log10(p value) on the y-axis. **B.** Heatmap representing the expression levels of selected genes in control and Long COVID-19 samples. Genes with higher expression in long COVID-19 samples are shown in warmer colors, whereas genes with higher expression in control samples are shown in cooler colors.

##### 3.3.1.3 Functional enrichment for the long COVID-19-only (LC-only) group

The functional enrichment of DEGs was performed to identify overrepresented biological functions, pathways, and cellular components, as shown by the KEGG pathway and Gene Ontology (GO) analyses. The enriched KEGG pathways are presented in **Figure 5A**. The most significant pathway, “Transcriptional misregulation in cancer,” is positioned at the top of the GeneRatio scale, with a large count and red color indicating a low adjusted p value. Other notably enriched pathways included microRNAs in cancer, NF-kappa B signalling, IL-17 signalling, and apoptosis across multiple species. Among these pathways, apoptosis in multiple species is the most statistically significant, despite having the fewest differentially expressed genes, making it a unique pathway for the long COVID-19 group.

**Figure 5:**
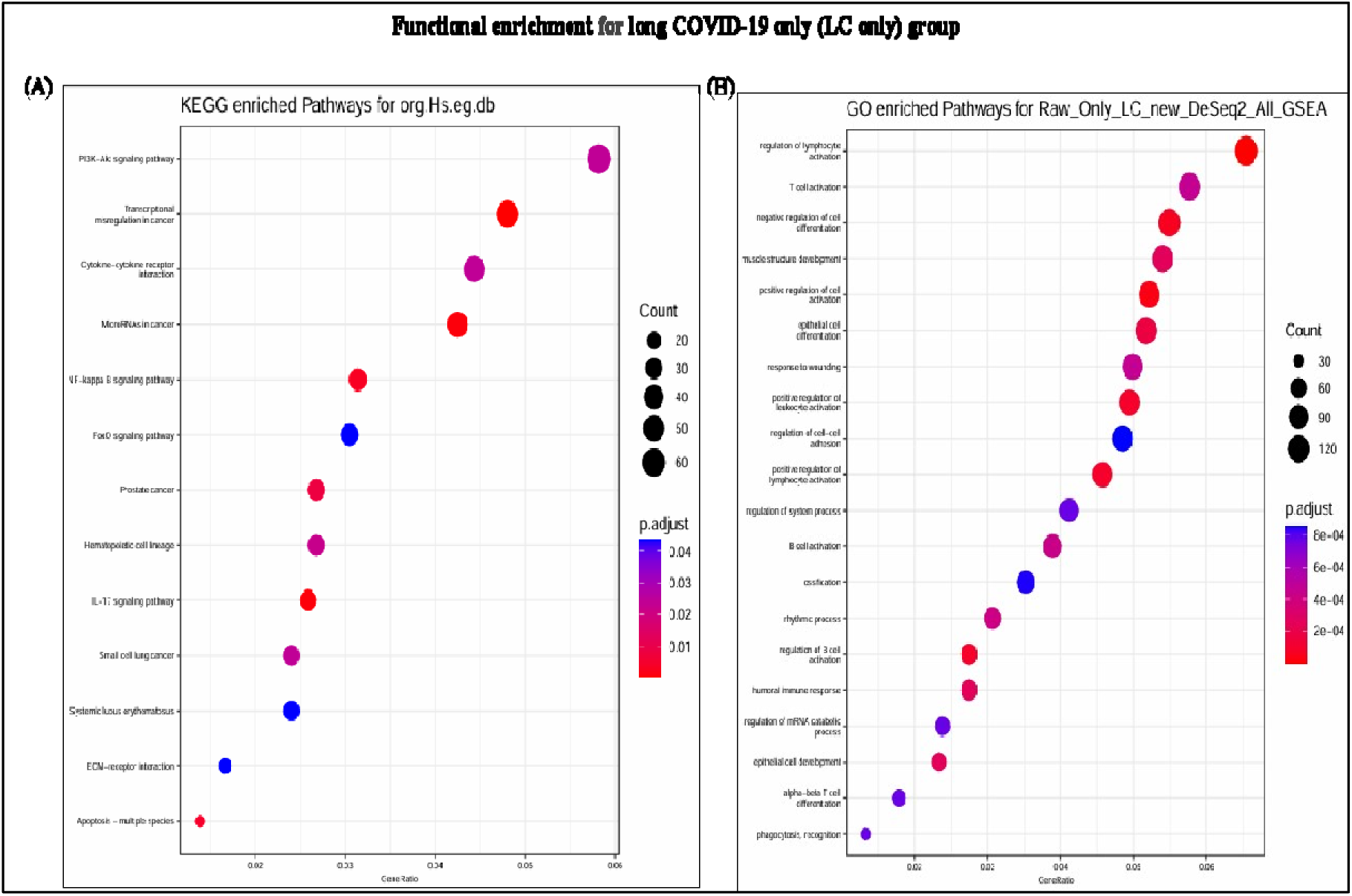
Functional enrichment of differentially expressed genes (DEGs) in Long COVID-19 (LC) patients only; **A.** KEGG pathway enrichment analysis. This dot plot shows the enriched KEGG pathways in the long COVID-19 group. The most significantly enriched pathway, “Transcriptional misregulation in cancer,” is highlighted with a large red dot, indicating high significance and a high gene ratio. Other notable pathways include microRNAs in cancer, NF-kappa B signalling, IL-17 signalling, and apoptosis across multiple species, with varying levels of gene involvement and significance. **B.** Gene Ontology (GO) biological process enrichment analysis. This dot plot displays the top enriched biological processes for the DEGs in long COVID-19 samples, including the regulation of lymphocyte activation, positive regulation of cell activation, B-cell activation, and other immune-related processes. The size of the dots represents the number of DEGs in each process, whereas the color intensity indicates the statistical significance of the enrichment.

The most significant Gene Ontology (GO) terms **(Figure 5B)** included regulation of lymphocyte activation, negative regulation of cell differentiation, positive regulation of cell activation, positive regulation of leukocyte activation, positive regulation of lymphocyte activation, and regulation of B-cell activation. These terms highlight the immune-related processes significantly enriched in the long COVID-19 samples.

#### 3.3.2 Healthy vs long COVID-19 brain fog (LCBF) group

##### 3.3.2.1 Top upregulated and downregulated DEGs

The total number of DEGs found in long COVID-19 (brain fog) patients was 207, of which 89 were upregulated and 118 were downregulated. The analysis of differentially expressed genes (DEGs) revealed significant upregulation of *EDNRB* and *SLC2A14,* with *EDNRB* showing the greatest fold change. On the other hand, several genes, such as *SMG1P1* and *CCL8*, were notably downregulated, with *SMG1P1* being the most significantly downregulated gene in the dataset. *NRCAM*, which we found among the exclusive genes, was also downregulated. These findings suggest a distinct molecular profile in the context of LCBF, as shown in **Table 3**.

**Table 3:** Top upregulated and downregulated DEGs in the Long COVID-19 brain fog (LCBF) group.

| UPREGULATED GENES |  |  | DOWNREGULATED GENES |  |  |
| --- | --- | --- | --- | --- | --- |
| Gene Symbol | log<sub>2</sub>FC | P-Value | Gene Symbol | log<sub>2</sub>FC | P-Value |
| *EDNRB* | 5.256 | < 0.001 | *SMG1P1* | -6.725 | < 0.000001 |
| *SLC2A14* | 4.532 | < 0.001 | *CCL8* | -5.203 | < 0.001 |
| *KRT17P1* | 4.484 | 0.047 | *NEBL* | -5.142 | 0.001 |
| *EDN2* | 4.390 | 0.007 | *AK4P3* | -4.729 | < 0.001 |
| *RBAK-RBAKDN* | 4.070 | 0.011 | *OR3A2* | -4.411 | < 0.001 |
| *None* | 4.041 | 0.038 | *PSMB11* | -4.406 | < 0.001 |
| *SIAH3* | 3.998 | 0.023 | *HSFX2* | -4.158 | 0.019 |
| *None* | 3.935 | 0.001 | *LINC02555* | -3.842 | 0.004 |
| *FAM47E-STBD1* | 3.925 | 0.046 | *CDC26P1* | -3.788 | 0.031 |
| *ANAPC1P4* | 3.844 | 0.001 | *None* | -3.778 | 0.046 |
| *None* | 3.647 | < 0.001 | *ELF2P1* | -3.778 | 0.024 |
| *CCL24* | 3.619 | 0.015 | *None* | -3.754 | 0.021 |
| *None* | 3.615 | 0.039 | *CROCCP2* | -3.743 | 0.044 |
| *H1-6* | 3.493 | 0.009 | *CCDC168* | -3.620 | 0.025 |
| *LINC02324* | 3.470 | 0.023 | *None* | -3.555 | 0.033 |
| *GSC* | 3.469 | < 0.001 | *None* | -3.542 | < 0.001 |
| *ZSCAN10* | 3.447 | 0.013 | *NRCAM* | -3.534 | < 0.000001 |
| *INHBA-AS1* | 3.390 | 0.003 | *HNRNPA1P16* | -3.479 | 0.019 |
| *NUAK1* | 3.375 | 0.001 | *CCNA1* | -3.464 | 0.013 |
| *SGCD* | 3.351 | 0.016 | *DSCAML1* | -3.436 | 0.021 |

##### 3.3.2.2 Volcano plot and heatmap for the long COVID-19 brain fog (LCBF) group

The volcano plot in **Figure 6A** shows differential gene expression between the LCBF and control samples. Significant genes such as *SMG1P1, NRCAM,* and *DHRS9* were downregulated, whereas *SLC2A14* was upregulated in the LCBF group. These genes are highlighted with distinct p values and fold changes, indicating their potential roles in the molecular mechanisms underlying LCBF. The heatmap **(Figure 6B)** further supports these findings by visualizing the log2-fold change values for the genes of interest across different samples. The LCBF and control groups presented different expression levels of *SMG1P1, NRCAM,* and *DHRS9*; these genes were downregulated.

**Figure 6:**
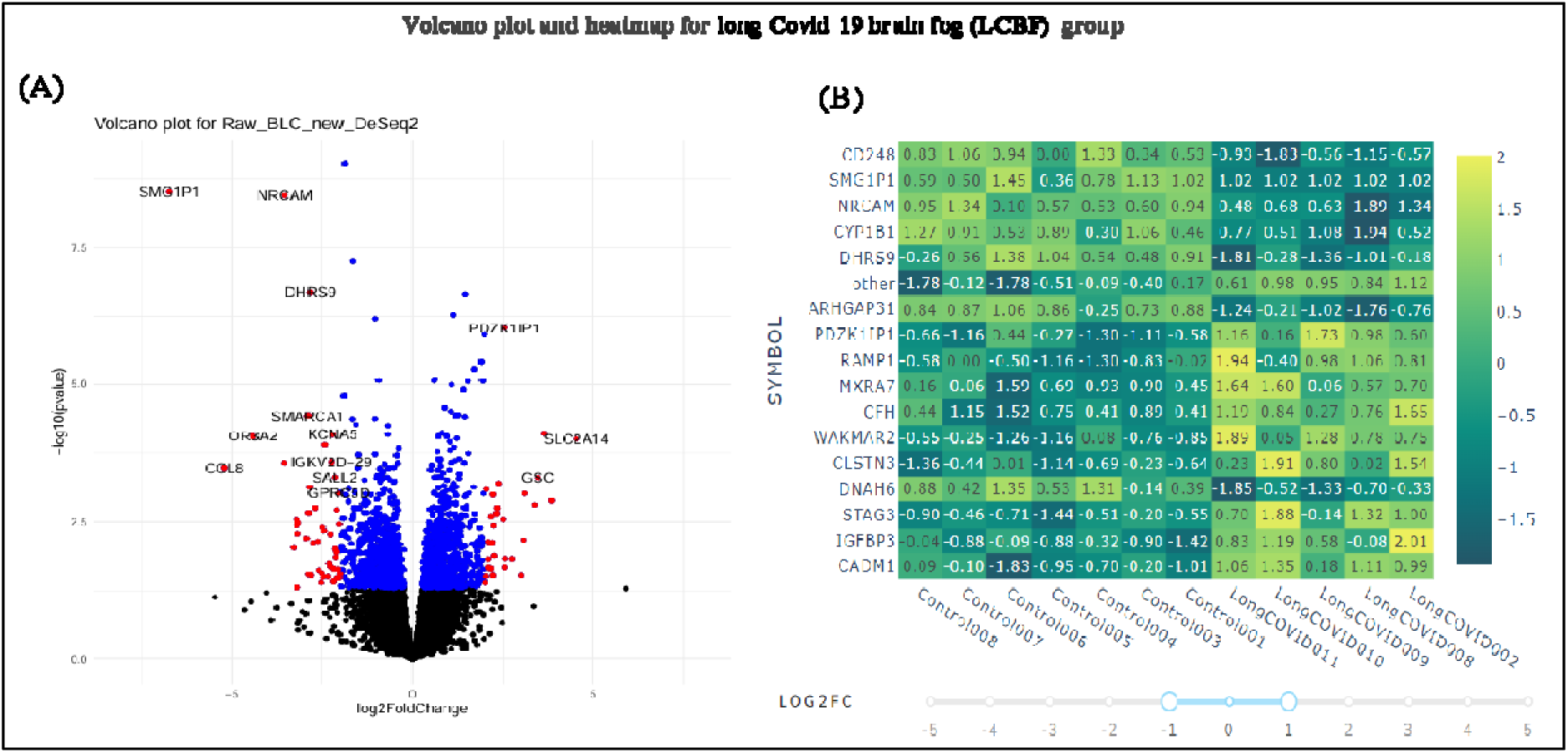
Differential gene expression analysis across the control and Long COVID-19 brain fog (LCBF) groups. **A.** Volcano plot showing the differential gene expression between control and long COVID-19 brain fog samples. The upregulated genes are displayed on the right, and the downregulated genes are on the left, with significance indicated by the -log10(p value) on the y-axis. **B.** Heatmap representing the expression levels of selected genes in control and Long COVID-19 brain fog samples. Genes with higher expression in long COVID-19 brain fog samples are shown in warmer colours, whereas genes with higher expression in control samples are shown in cooler colours.

##### 3.3.1.3 Functional enrichment in the long COVID-19 brain fog (LCBF) group

Owing to sub-optimal results from the T-Bio platform, we utilized ENRICHR for KEGG pathway analysis. **Figure 7A** shows that the KEGG pathway analysis revealed that LCBF affects several crucial pathways, including hypertrophic cardiomyopathy, dilated cardiomyopathy, relaxin signalling, calcium signalling, viral myocarditis, and renin secretion. **In Figure 7B**, the Gene Ontology molecular function analysis highlights “transmembrane signalling receptor activity” as particularly significant, as indicated by its large dot size, low p.adjust value, and relevance to the condition of LCBF.

**Figure 7:**
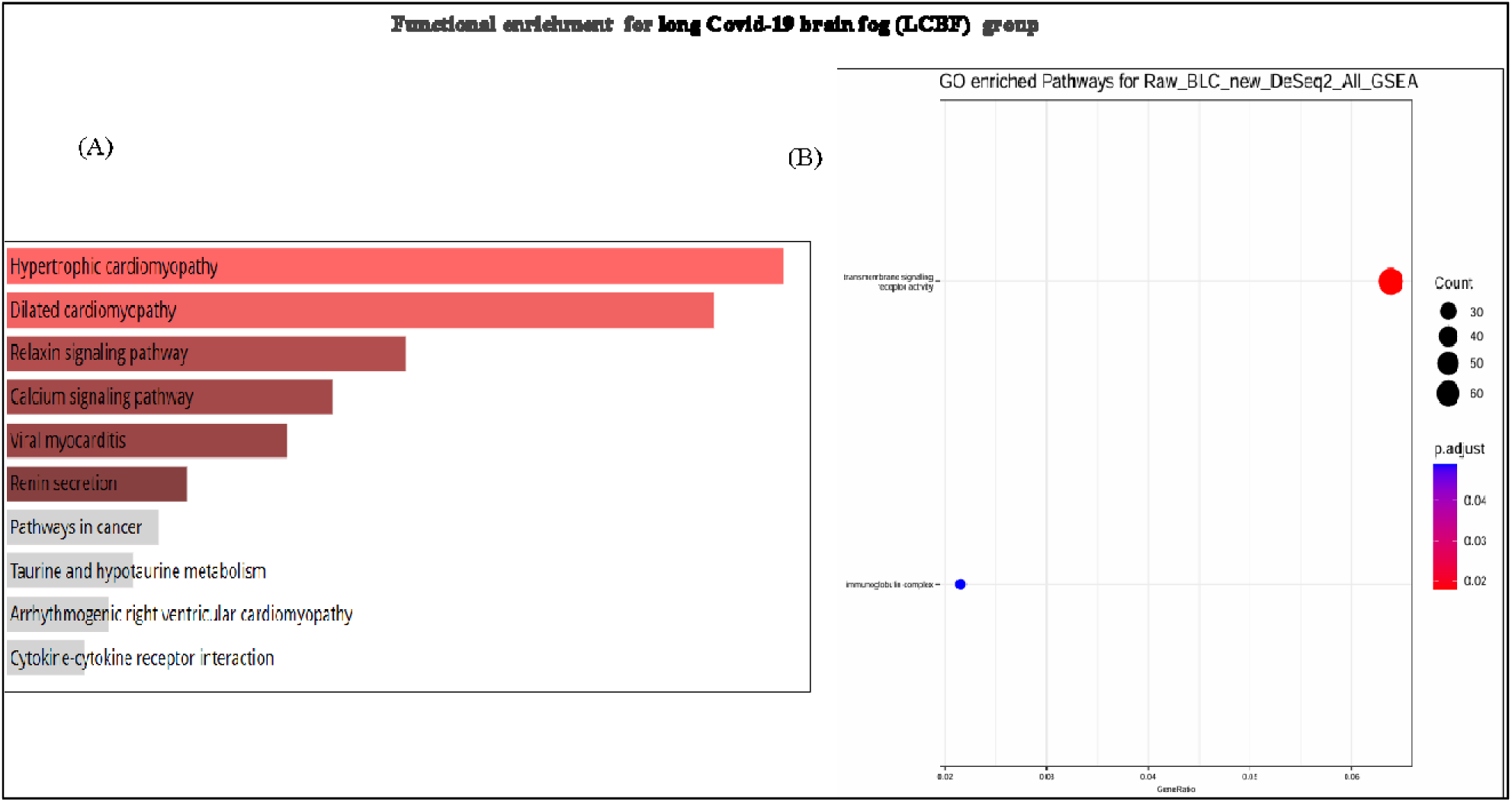
KEGG and Gene Ontology (GO) pathway enrichment analysis for Long COVID-19 brain fog (LCBF) samples. **A.** KEGG pathway enrichment analysis: The bar plot in Panel A displays the top KEGG pathways significantly enriched in the long COVID-19 brain fog group. Prominent pathways include “hypertrophic cardiomyopathy,” “dilated cardiomyopathy,” and “relaxin signalling pathway.” These pathways are closely linked to cardiovascular dysfunction, and their strong importance highlights their potential relevance to the pathophysiology of LCBF. **B.** GO pathway enrichment analysis: The dot plot in Panel B shows the most significantly enriched GO terms for biological processes in long COVID-19 brain fog samples. “Transmembrane signalling receptor activity” emerged as the most significant biological process, with a high gene ratio, suggesting its crucial involvement in receptor signalling mechanisms associated with LCBF.

#### 3.3.3 Healthy vs Convalescent (Con)

##### 3.3.3.1 Top upregulated and downregulated DEGs

The total number of DEGs found in Convalescent patients was 650, of which 332 were upregulated genes and 318 were downregulated genes. **Table 4** shows that HBG1 was one of the genes with upregulated expression in convalescent patients, whereas GPRC5A and RASD2 had the most downregulated expression.

**Table 4:**
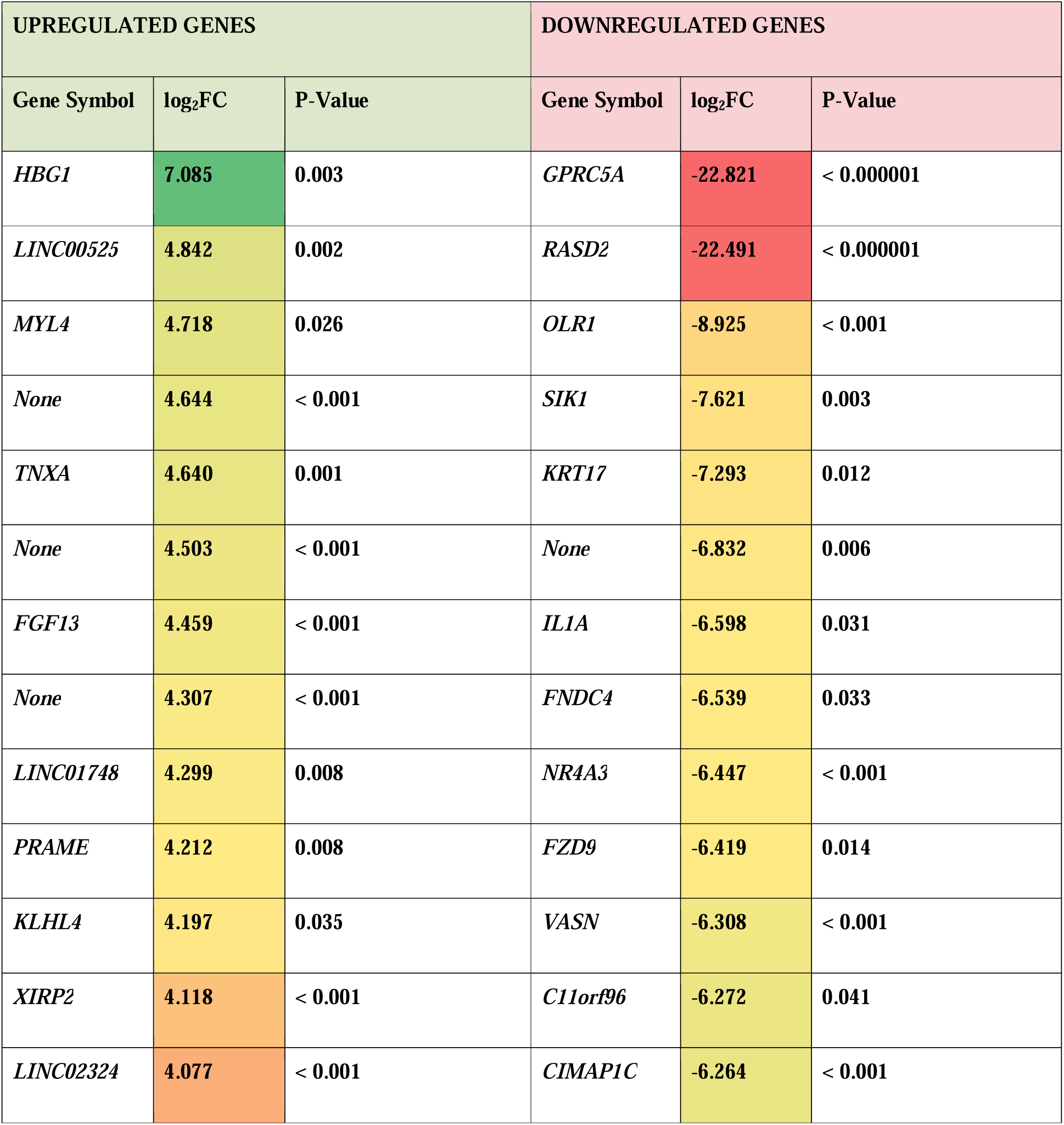
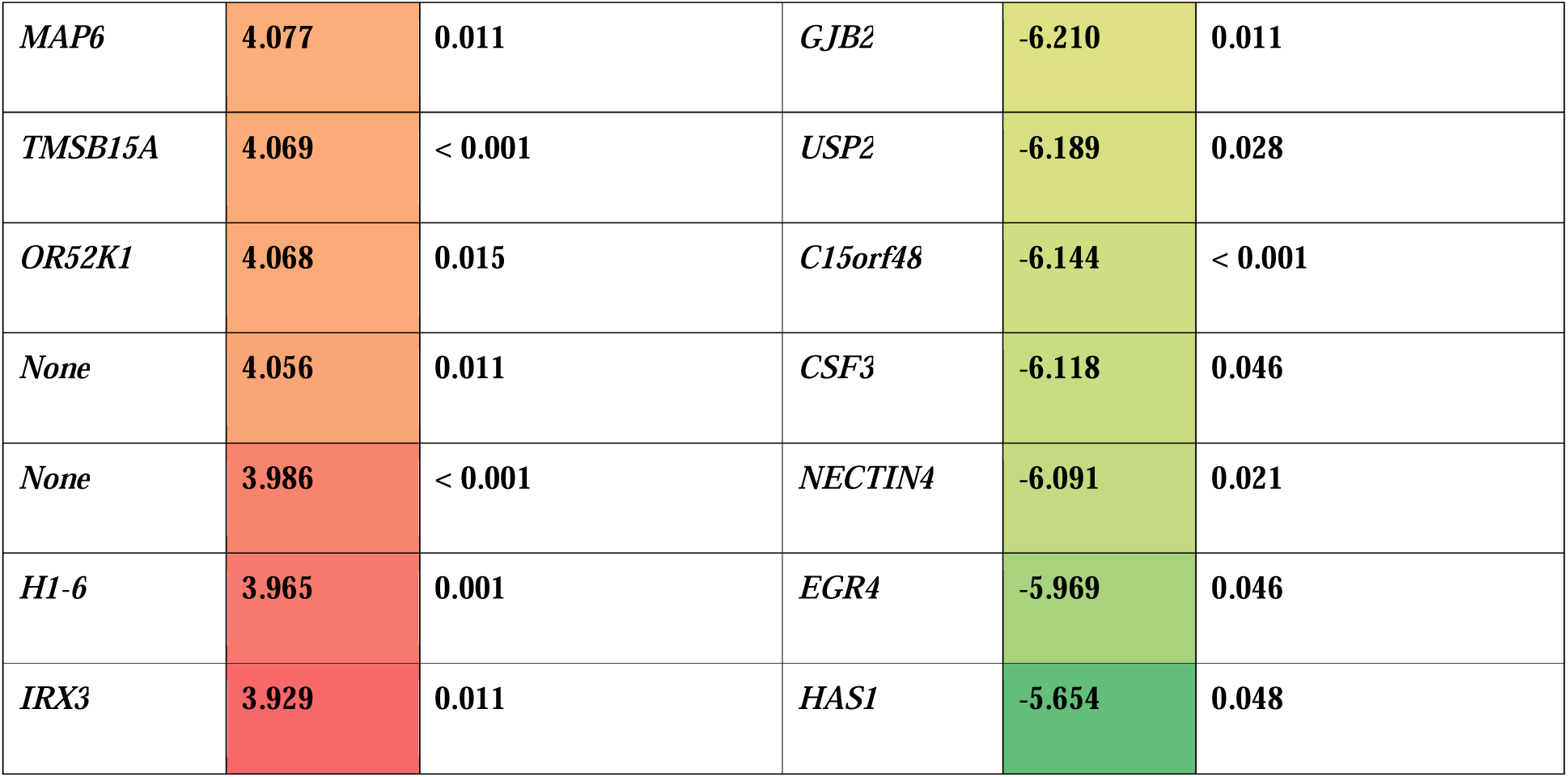
Top upregulated and downregulated DEGs in the Convalescent (Con) group.

##### 3.3.3.2 Volcano plot and heatmap for the Convalescent (Con) group

The volcano plot shows that *GPRC5A* is significantly downregulated (**Figure 8A).** The heatmap (**Figure 8B)** shows that most genes in the convalescent group seem to be upregulated and that those in the control group appear to be downregulated, except for the genes *GPRC5A* and *RASD2,* which are overexpressed but are underexpressed in the convalescent group in the samples.

**Figure 8:**
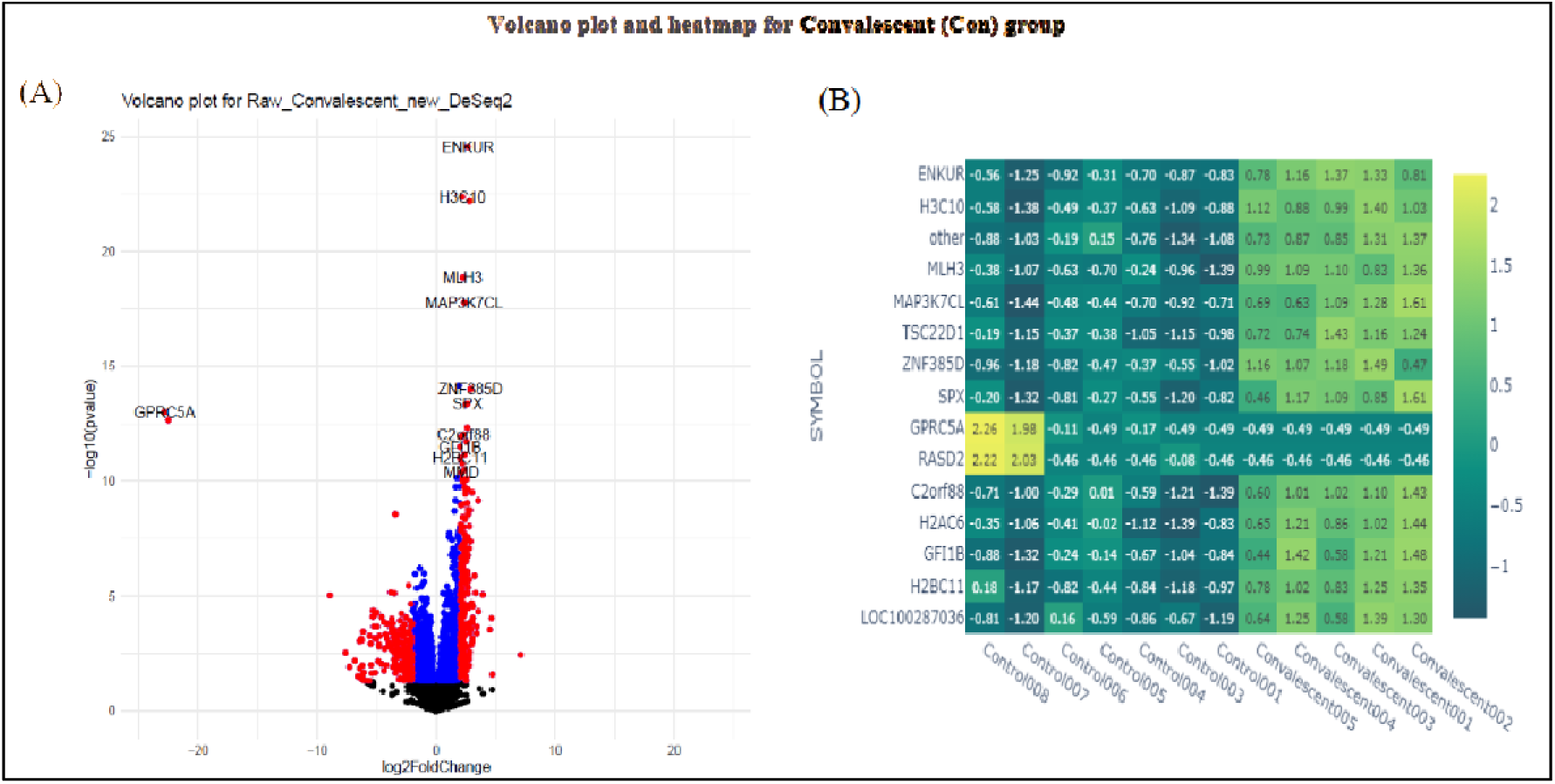
Differential gene expression analysis across the control and Convalescent (Con) groups. **A.** Volcano plot showing the differential gene expression between the control and Convalescent samples. The upregulated genes are displayed on the right, and the downregulated genes are on the left, with significance indicated by the - log10(p value) on the y-axis. **B.** Heatmap representing the expression levels of selected genes in control and Convalescent samples. Genes with higher expression in Convalescent samples are shown in warmer colors, whereas genes with higher expression in control samples are shown in cooler colors.

##### 3.3.3.3 Functional enrichment for the Convalescent (Con) group

KEGG pathway enrichment analysis **(Figure 9A)** revealed significant enrichment of several pathways in the convalescent group, including “Transcriptional misregulation in cancer,” “Neuroactive ligand□receptor interaction,” and “Hematopoietic cell lineage.” These pathways, particularly those related to transcriptional misregulation in cancer, stand out with low p-adjusted values, indicating their potential involvement in the convalescent group’s biological response. GO pathway enrichment analysis **(Figure 9B)** revealed that biological processes such as “vasculature development,” “blood vessel development,” and “wound healing” were highly enriched in the convalescent group. These processes, with low p-adjust values, suggest that the convalescent samples might exhibit enhanced wound-healing processes compared with the controls, reflecting their postrecovery stage.

**Figure 9:**
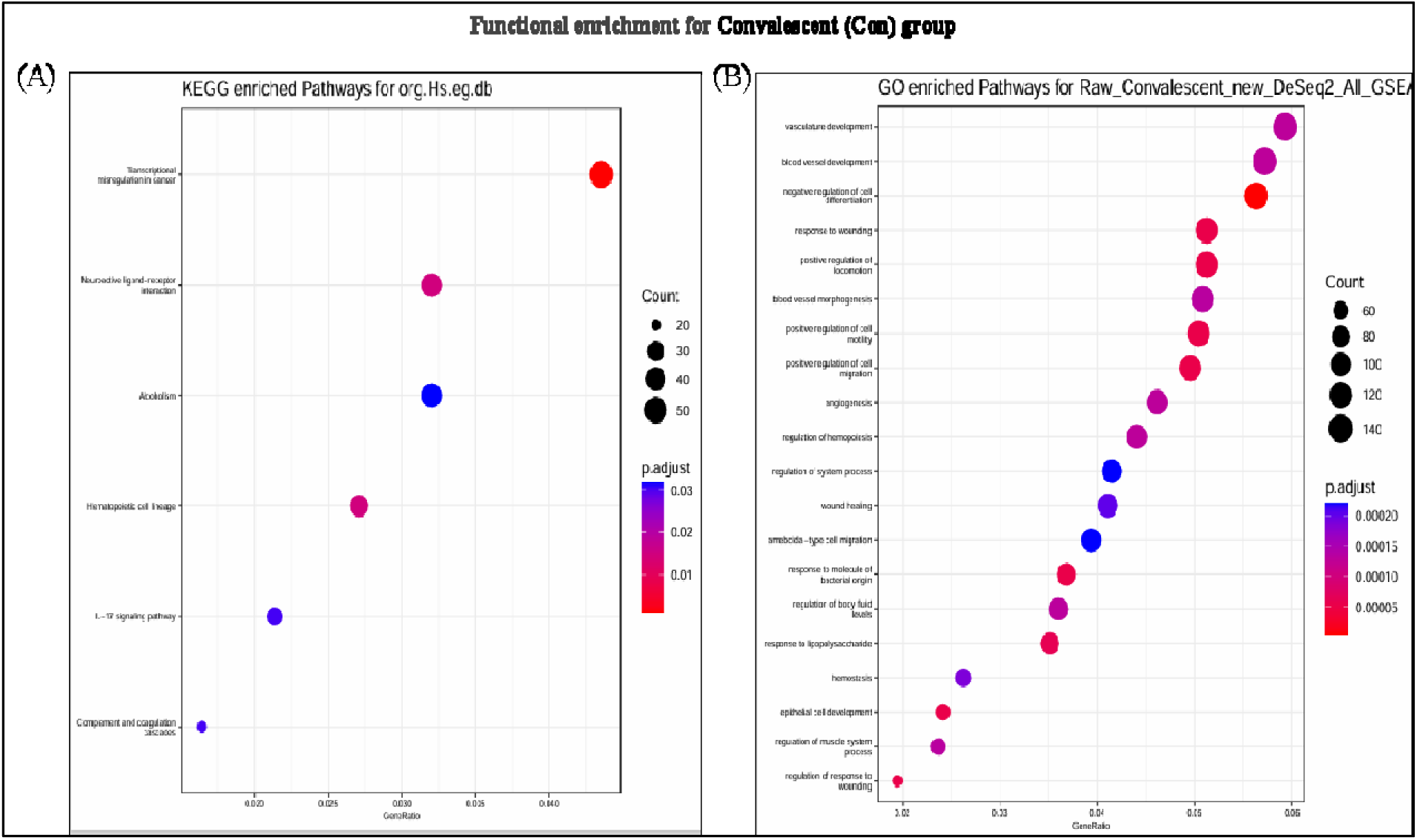
KEGG and Gene Ontology (GO) pathway enrichment analysis for the Convalescent (Con) group. **A.** KEGG pathway enrichment analysis: Dot plot showing the significant KEGG pathways enriched in the Convalescent group, highlighting pathways such as “Transcriptional misregulation in cancer,” “Neuroactive ligandLreceptor interaction,” and “Hematopoietic cell lineage.” The size and color of the dots indicate the gene ratio and statistical significance, with red and purple dots representing the most significant pathways. **B.** GO pathway enrichment analysis: Dot plot showing the most significantly enriched biological processes in the convalescent group. Key processes such as “vasculature development,” “blood vessel development,” and “wound healing” were enriched, with the size and color of the dots indicating the gene ratio and p.adjust values.

### 3.4 Comparative analysis

**Figure 10** displays a Venn diagram of the DEGs across three pairwise comparisons: healthy versus convalescent (H vs CON), healthy versus long COVID-19 only (H vs LC), and healthy versus long COVID-19 with brain fog (H vs BLC). In the H vs CON comparison, 313 DEGs (47.1%) were unique to convalescent samples, 55 DEGs (8.3%) were unique to the H vs LC group, and 107 DEGs (16.1%) were unique to the H vs BLC group. There were 154 DEGs (23.2%) shared between H vs CON and H vs LC, 20 DEGs (3.0%) shared between H vs CON and H vs BLC, and 4 DEGs (0.6%) shared between H vs LC and H vs BLC. Notably, 11 DEGs (1.7%) were common to all three comparisons **(see Table 4)**.

**Figure 10:**
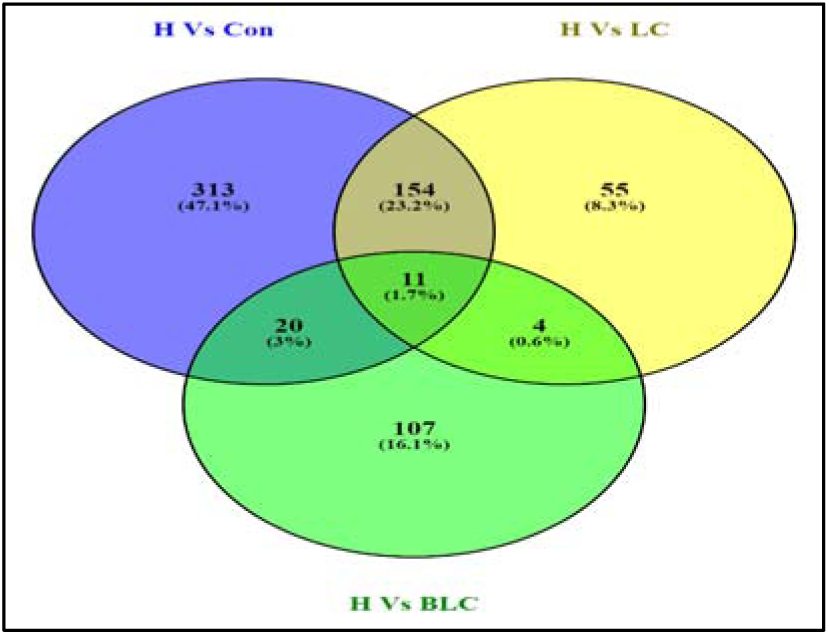
Venn diagram illustrating the distribution and overlap of differentially expressed genes (DEGs) across three groups: Healthy vs Convalescent. (H vs. CON), Healthy vs Long COVID-19 (H vs. LC), and Healthy vs Long COVID-19 with Brain Fog (H vs. BLC). The diagram shows unique and overlapping DEGs between these groups, with 11 common DEGs shared among all three comparisons.

### 3.5 Pathway and GO analysis of exclusive DEGs in the LCBF group via Enrichr

#### 3.5.1 KEGG pathway

The KEGG pathway analysis **(Figure 11A)** for the exclusive genes in the LCBF group highlighted the “relaxin signalling pathway” as the most statistically significant. NOS2 was found to be involved in several key pathways, including relaxin signalling and arginine biosynthesis **(Figure 11B).**

**Figure 11:**
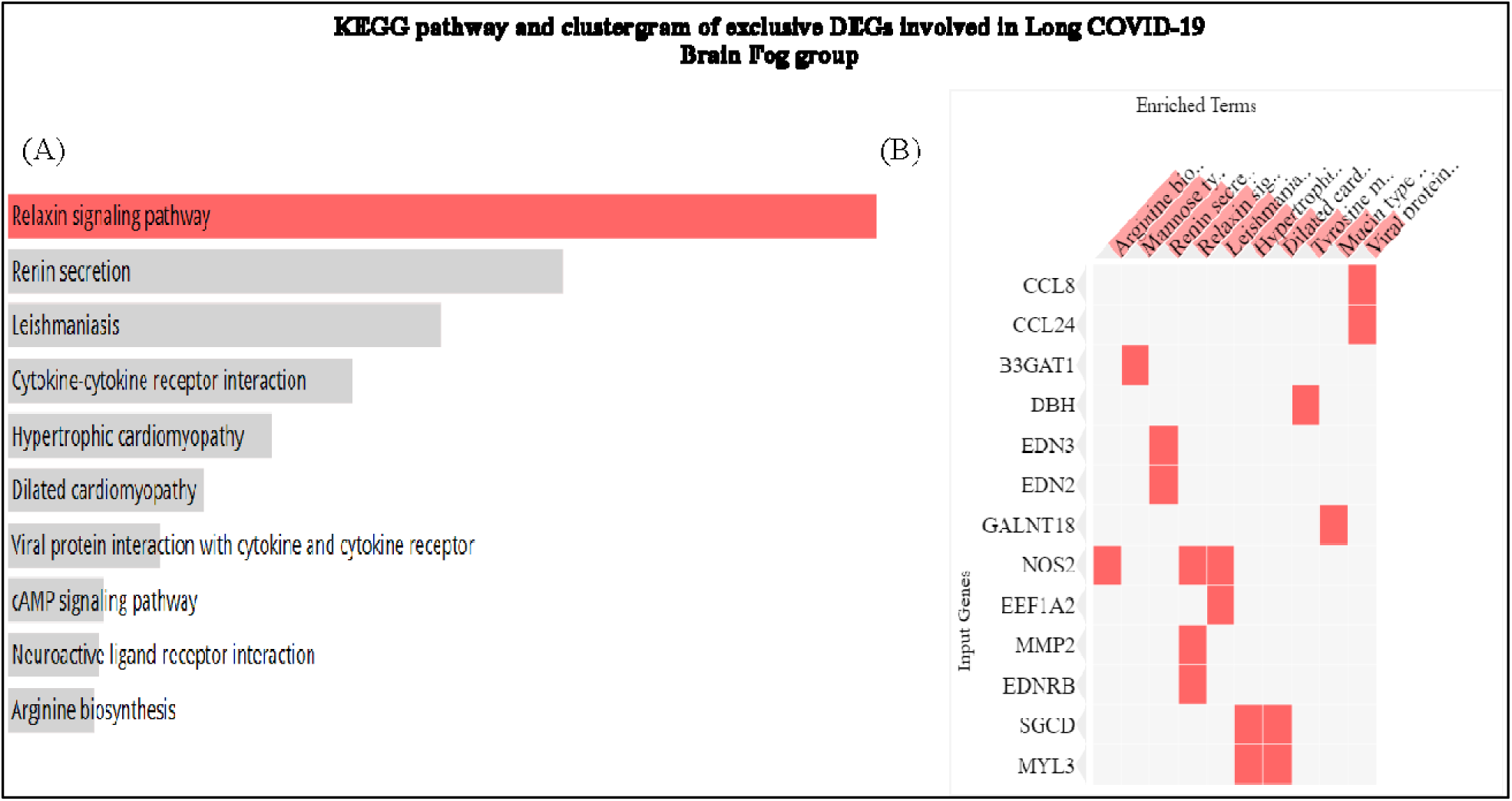
A. KEGG pathway enrichment analysis of exclusive genes in the LCBF group. Significant pathways, such as the “relaxin signalling pathway” and “arginine biosynthesis”, are shown. **B.** Clustergram illustrating the exclusive genes involved in the significant KEGG pathways in the LCBF group.

#### 3.5.2 Gene Ontology

Gene Ontology (GO) analysis of biological processes, as depicted in **Figure 12A**, revealed that vascular-associated smooth muscle contraction, vasoconstriction, and phasic smooth muscle contraction were the most significantly enriched terms. The genes *EDNRB, EDN2,* and *EDN3* were closely associated with these processes, as shown in **Figure 12B**, suggesting a potential link between vascular dysfunction and cognitive symptoms in the LCBF.

**Figure 12:**
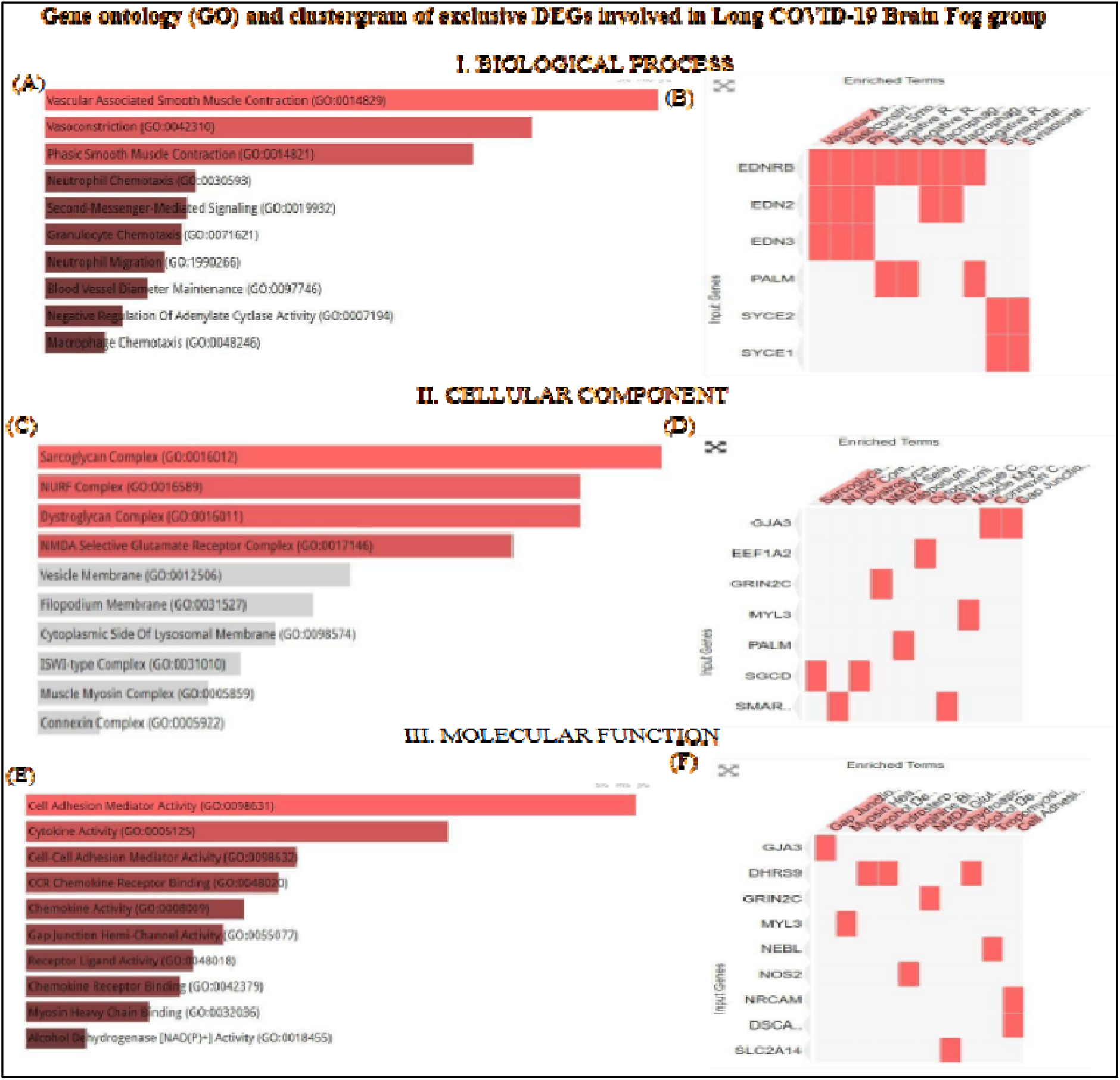
Gene Ontology (GO) and clustergram of exclusive DEGs involved in the long COVID-19 brain fog (LCBF) group. **A.** Bar graph showing the most statistically significant biological processes in the LCBF group. Vascular-associated smooth muscle contraction and vasoconstriction were the key terms enriched. **B.** Clustergram of exclusive genes involved in the biological processes in the LCBF group. **C.** Bar graph of the cellular components associated with the LCBF group. Significant components include the Sarcoglycan Complex and the NMDA Selective Glutamate Receptor Complex. **D.** Clustergram of exclusive genes associated with cellular components in the LCBF group. **E.** Bar graph showing the most significant molecular functions in the LCBF group, including cell adhesion mediator activity and cytokine activity. **F.** Clustergram of exclusive genes involved in the molecular functions in the LCBF group.

Furthermore, cellular component analysis, illustrated in **Figure 12C**, identified the sarcoglycan complex, NURF complex, Dystroglycan complex, and NMDA selective glutamate receptor complex as critical components in the LCBF group. GRIN2C, associated with the NMDA glutamate receptor complex, is crucial for synaptic plasticity and cognitive functions, as depicted in **Figure 12D**.

Finally, molecular function analysis, presented in **Figure 12E**, revealed that cell adhesion mediator activity, cytokine activity, and cell–cell adhesion mediator activity were the most significant functions. Genes such as *DHRS9* and *NRCAM* are involved in these pathways, as highlighted in **Figure 12F**, underscoring their relevance to cognitive disruptions in long-term COVID-19 patients.

Distinct transcriptomic signatures were identified exclusively in brain fog patients, notably upregulated genes including GRIN2C (an NMDA receptor subunit), EDNRB (associated with endothelin signaling), and NOS2 (inducible nitric oxide synthase). KEGG and GO analyses revealed significant enrichment in pathways including calcium signaling, cardiovascular dysfunction (e.g., hypertrophic and dilated cardiomyopathy), and relaxin signaling, implicating these pathways in cognitive impairment.

## 4. DISCUSSION

This study revealed that long-term COVID-19 patients with brain fog possess a distinct peripheral gene expression profile, distinguishing them from those without cognitive issues and those who have fully recovered (convalescent individuals). This was evident from the principal component analysis (PCA), where the brain fog group clustered separately, suggesting a coherent molecular signature associated with cognitive impairment. Notably, although all long-term COVID-19 patients had a common infection history, those with brain fog presented specific transcriptomic differences, indicating ongoing perturbations related to their cognitive symptoms. Our findings are consistent with those of Greene *et al*. (2024), who reported unique immune profiles in long-term COVID-19 patients with cognitive complaints.

Having established the distinct molecular signature associated with cognitive impairment, we now delve into specific gene expression, beginning with the long COVID-19-only group (without brain fog). Transcriptomic analysis revealed significant upregulation of *IGHV7-4-1*, a gene involved in B-cell responses. This gene is also enriched in MDA5+ dermatomyositis (DM) and COVID-19, which involve B-cell activation, although they differ in their type I interferon signalling profiles (Ye *et al*., 2022). Skewed expression of the *IGHV, IGLV,* and *IGKV* genes, particularly the *IGHV7-4-1* and *IGLV3-27* combination, has been observed in asymptomatic COVID-19 patients (Ma *et al*., 2022). The link between MDA5+ DM and rapidly worsening interstitial lung disease (ILD) might account for the high levels of *IGHV7-4-1* gene expression in COVID-19 patients. Interestingly, “regulation of B-cell activation” was among the most statistically significant GO terms, suggesting a potential role in antibody-based therapeutic strategies. In long COVID-19, considerable downregulation of *CHMP1B-AS1* was observed. *CHMP1B-AS1* is a long noncoding RNA antisense to the *CHMP1B* gene, which encodes an ESCRT-III endosomal sorting protein. ESCRT complexes are essential for endocytic recycling and neuronal homeostasis (Lee and Gao, 2012). Dysregulated *CHMP1B-AS1* could thus impact CHMP1B/ESCRT function. Moreover, many enveloped viruses exploit the ESCRT pathway for budding (Ju *et al*., 2021). In summary, *CHMP1B-AS1* is likely related to ESCRT-dependent membrane dynamics, with indirect implications for viral infection cycles and neural health. In all long COVID-19-only samples, we also observed a reduction in *OLR1*, a gene previously linked to vascular protection. Its decrease suggests potential vascular and inflammatory complications post-COVID-19, as observed in conditions such as osteoarthritis (Ohno *et al*., 2021; Tugarov *et al*.,2023). Notably, reduced *OLR1* expression is associated with an increased risk of Alzheimer’s disease (Serpente *et al*., 2011), raising concerns about cognitive sequelae in COVID-19 patients. Pathway analysis revealed that “transcriptional misregulation in cancer” was a significantly enriched pathway, suggesting that cancer-like dysregulation of gene expression may underlie persistent inflammation and immune dysfunction (Saini & Aneja, 2021; Kavanagh, 2022). Other enriched pathways included NF-κB, IL-17, microRNAs in cancer, and apoptosis, indicating complex immune□inflammatory interactions.

In contrast, the convalescent group presented a broader range of differentially expressed genes (650 DEGs), reflecting an ongoing recovery process. *HBG1* (Haemoglobin Subunit Gamma 1) stands out among the upregulated genes. Its elevated expression may reflect ongoing respiratory adaptation or residual inflammation (Majeed & Shajar, 2020). Among the downregulated genes, *GPRC5A* and *RASD2* were of particular interest. *GPRC5A,* which is highly expressed in lung tissues, functions as a tumor suppressor in lung cancer and an oncogene in other cancers (Dai *et al*., 2021; González *et al*., 2023; Jiang *et al*., 2018). Its suppression suggests possible implications for postviral lung pathology. *RASD2*, a Ras GTPase family member, is involved in mood regulation. Its downregulation is associated with depression- and schizophrenia-like behaviours (Cheng *et al*., 2022; Vitucci *et al*., 2015) and may correlate with neuropsychiatric symptoms, including the cognitive disturbances observed in convalescent COVID-19 patients. Pathway enrichment in this group highlighted biological processes related to vasculature development, reflecting vascular remodelling. For instance, Patel *et al*. (2022) reported that angiogenesis□related biomarkers (e.g., angiopoietin□1, P□selectin) were markedly elevated in Long COVID-19 patients relative to controls. These findings suggest that ongoing vascular transformation and new vessel formation occur in response to endothelial injury (Patel *et al*., 2022). The enrichment of wound healing-related biological processes suggested that SARS-CoV-2 infection induces a tissue repair and regeneration response, even during the early phase of infection. This activation reflects the host’s attempt to counteract virus-induced tissue damage (Lieberman *et al*., 2020).

Focusing on the LCBF group, we identified 207 DEGs, 89 of which were upregulated and 118 of which were downregulated. Among the upregulated genes, *SLC2A14,* encoding the glucose transporter *GLUT14,* was notable. Its upregulation implies altered glucose transport and metabolism in the brain, potentially contributing to cognitive dysfunction (Shaghaghi *et al*., 2016). Conversely, *SMG1P1* downregulation suggests impaired gene expression stability, possibly exacerbating synucleinopathies such as Parkinson’s disease (Henderson-Smith *et al*., 2013). **Enrichr** pathway analysis revealed enrichment in cardiac-related pathways such as hypertrophic cardiomyopathy, dilated cardiomyopathy, and viral myocarditis, supporting emerging evidence of cardiovascular involvement in long COVID-19-related brain fog (Chatys-Bogacka *et al*., 2024; Mailian & Kolomiets, 2019; Schrank, Barrington, & Stutzmann, 2019). Transmembrane signalling receptors include many immune and neural receptors that span the cell membrane (e.g., cytokine receptors, TLRs, and neurotransmitter receptors). Abnormal signalling through these receptors has been implicated in long COVID-19 brain fog. For example, Fontes□Dantas *et al*. (2023) demonstrated that the SARS-CoV-2 spike protein activates Toll□like receptor 4 (TLR4, a transmembrane innate immune receptor) to induce synaptic loss and persistent memory impairment in mice. Similarly, chronic post□COVID-19 neuroinflammation involves cytokine cascades that dysregulate microglia and cause cognitive symptoms (Low, Low, and Akrami, 2023). Fesharaki□Zadeh *et al*. (2023) reported that elevated kynurenic acid after infection blocks NMDA□ and nicotinic□α7 neurotransmitter receptors (both transmembrane), leading to prefrontal cortical deficits characteristic of brain fog. These studies link aberrant transmembrane receptor signalling (immune and neurotransmitter receptors) to sustained inflammation and cognitive dysfunction in long COVID.

Venn analysis revealed 11 DEGs **(Table 5)** common across all comparisons (long COVID-19, brain fog, and convalescence), indicating a shared molecular response to SARS-CoV-2. However, the identification of 107 genes exclusive to brain fog highlights its distinct molecular signature.

**Table 5:**
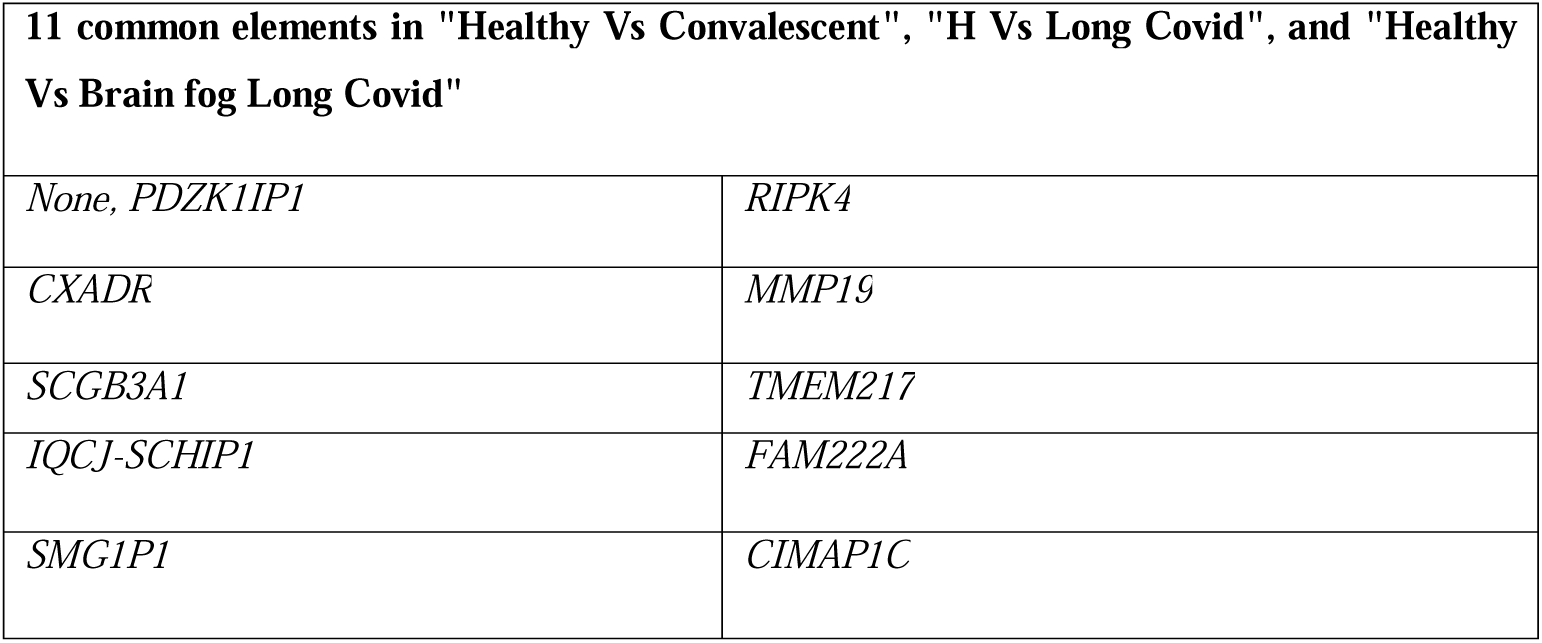
Common DEGs in “Healthy vs Convalescent”, “Healthy vs Long Covid”, and “Healthy vs Long Covid-19 with brain fog”.

The NMDA receptor subunit gene *GRIN2C* was one of the exclusive DEGs in the brain fog group. NMDARs are critical for synaptic plasticity and cognition. Altered *GRIN2C* expression may disrupt cortical excitatory–inhibitory balance, impairing cognitive functions (Gupta *et al*., 2016). Lower *GRIN2C* levels have been associated with schizophrenia and late-onset Alzheimer’s disease (Rubino *et al*., 2025). Given the role of *GRIN2C* in synaptic plasticity and its exclusivity in our dataset suggests a possible link to the cognitive symptoms observed in long-term COVID-19 patients, although further studies are needed to establish a direct causal relationship.

Several genes implicated in neurovascular and inflammatory processes were also upregulated. *EDNRB*, encoding the endothelin-B (ET_B) receptor, was the most highly upregulated gene. Endothelin signalling is crucial for cerebrovascular tone and BBB integrity (Alcendor, 2020). Excessive endothelin-1 (ET-1) activity through its receptors (ET_A and ET_B) activity can promote vasoconstriction, chronic hypoperfusion, and inflammation in the brain (Alcendor, 2020). The upregulation of *EDNRB* in astrocytes may contribute to reactive astrocyte formation and cognitive deficits (Koyama, 2021; Michinaga *et al*., 2023).

Markers of **innate immune activation** were also prominent among the brain fog-specific differentially expressed genes. Similarly, the exclusive DEG *NOS2*, encoding inducible nitric oxide synthase (iNOS), suggests ongoing neuroinflammation. Excessive nitric oxide production during viral infections is neurotoxic and may impair synaptic function (Cespuglio *et al*., 2021) In viral infections of the CNS, excessive iNOS-derived nitric oxide can be neurotoxic; for example, in a mouse coronavirus encephalitis model, mice lacking NOS2 had reduced neuronal apoptosis and better survival, implicating iNOS in virus-induced neuronal injury (Chen & Lane, 2002). In the context of SARS-CoV-2, elevated cytokines and activated glia can drive a surge in iNOS and nitric oxide production in the brain (Cespuglio et al., 2021). While nitric oxide can have antiviral effects, an overproduction of NO and its reactive derivatives (like peroxynitrite) is deleterious to neural cells and synapses as it cause oxidative damage (Cespuglio et al., 2021). The relaxin signalling pathway, in which *NOS2* is involved, emerged among the top enriched pathways, indicating a potential counterregulatory mechanism attempting to restore vascular and neural homeostasis because relaxin is a hormone known to have anti-fibrotic and vasodilatory effects (Garvin, 2018; Chow *et al*., 2012). By upregulating eNOS and modulating iNOS activity, relaxin can enhance NO bioavailability, reduce oxidative stress, and promote BBB integrity (Baccari & Bani, 2008) (Hoos *et al*., 2014). **Clinically, targeting the NOS2 gene and the relaxin signaling pathway could offer therapeutic potential for alleviating brain fog in long COVID (Garvin, 2018).** For instance, pharmacological agents that enhance relaxin signaling or modulate iNOS activity may reduce neuroinflammation and oxidative stress, thereby improving cognitive function.

Other downregulated genes, such as *DHRS9* and *NRCAM,* further support the hypothesis of impaired immune regulation and compromised BBB integrity. *DHRS9* downregulation results in reduced regulatory macrophage activity and diminished retinoic acid production, exacerbating inflammation (Riquelme *et al*., 2017). When *NRCAM,* which is essential for axon guidance and BBB maintenance, is downregulated, it may exacerbate BBB permeability and allow infiltration of inflammatory cytokines into the CNS (Moy *et al*., 2009; Greene *et al*., 2024).

Pathway enrichment analysis revealed that long-term COVID-19 brain fog involves unique biological processes. Several **KEGG pathways, including calcium signalling and dilated cardiomyopathy,** were significantly enriched only in the brain fog group**. Calcium signalling pathway** enrichment indicates disturbances in cellular calcium homeostasis in the brains of patients with fog. Calcium signalling is fundamental for neuronal communication, muscle contraction, and virtually all cell types; thus, its dysregulation could have wide-ranging effects. Excessive intracellular Ca^2+^ levels in neurons can lead to excitotoxic damage. Recent neuropathological studies on COVID-19 have revealed that calcium regulation is involved in **Alzheimer’s disease-like changes**. Reiken *et al*. (2022) reported that the brains of COVID-19 patients exhibit hyperactivation of Ca^2+^-releasing channels (ryanodine receptor 2) due to oxidative stress, resulting in “leaky” calcium channels similar to those observed in Alzheimer’s disease. This calcium leak can impair synaptic function and induce cognitive deficits (Reiken *et al*. 2022). Our pathway results align with these findings, suggesting that long-term COVID-19 brain fog may involve chronic disruptions in Ca2+ signalling cascades. The enrichment of **dilated cardiomyopathy (DCM) pathway genes** in patients with brain fog could reflect ongoing cardiovascular stress or **autonomic dysregulation,** which is uniquely associated with cognitive symptoms. It is well established that cardiac insufficiency can lead to cognitive impairment owing to diminished cerebral perfusion and embolic phenomena (Iadecola *et al*., 2016). In patients with heart failure, chronic low cardiac output and hypotension result in cerebral hypoperfusion, which causes white matter damage, cortical atrophy, and cognitive decline over time (Wolters *et al*., 2017; Alosco *et al*., 2013). Similarly, persistent cardiac inflammation or autonomic abnormalities (such as postural orthostatic tachycardia syndrome) can limit blood flow to the brain, leading to fatigue and fog (Haloot *et al*., 2023).

Validation of transcriptomic biomarkers such as *NOS2, EDNRB, DHRS9, GRIN2C,* and *NRCAM*, alongside enriched pathways (calcium signalling, relaxin, endothelin, cardiomyopathy), all enriched in CNS tissue or on membranes requires integrated molecular and functional assays to enable clinical translation. Because these transcripts are poorly detected in blood, validation relies on surrogate readouts. *GRIN2C* and *NRCAM* can be tracked via CNS□derived exosomes or CSF proteomics (Pulliam *et al*., 2019) (Sun *et al.,* 2019) (Reinhold *et al*., 2023). Measuring CSF glutamate and glutamine levels via high performance liquid chromatography (HPLC) provided insights into NMDA receptor-related excitotoxicity, serving as an indirect method to assess *GRIN2C* activity (Madeira *et al*., 2018). It was found that long-COVID patients had a higher plasma glutamine/glutamate ratio than controls (López-Hernández *et al.,* 2023). More directly, Li *et al*. (2024) reported elevated CSF quinolinic acid (an NMDA-receptor agonist) in post-COVID patients, which correlated with other markers of neurodegeneration. PET, MRI/fMRI and EEG provide noninvasive brain biomarkers. FDG-PET often shows focal hypometabolism in long-COVID patients (Talkington *et al*., 2025). High-field fMRI connectivity correlates with cognitive impairment (Barnden *et al.,* 2023). EEG studies reveal persistent network abnormalities (e.g. reduced alpha power) after COVID (Y. Sun *et al.,* 2024). Given the enrichment of cardiomyopathy pathways, cardiac biomarkers and imaging should be considered. Echocardiography (echo) is a first-line tool: in COVID-19 it often reveals LV/RV dysfunction, strain abnormalities, or pericardial effusion (Crosier *et al.,* 2023). S100β and GFAP are established astroglial injury markers, while blood test like troponin-T indicates cardiomyocyte damage (Greene *et al.,* 2024) (Rêgo *et al.,* 2023). Greene *et al*. (2024) found that acute COVID-19 patients with cognitive impairment had significantly higher serum S100β than controls, implicating BBB leakage. Their long-COVID cohort (“brain fog” subgroup) also showed elevated GFAP, which is known to rise after cerebrovascular injury. Meanwhile, cardiac troponin-T is often elevated in COVID-19 with myocarditis or cardiomyopathy (Rêgo *et al.,* 2023). Thus, a peripheral profile of high S100β/GFAP (reflecting astrocyte/BBB injury) together with high troponin-T (reflecting COVID-related cardiac damage) would support the link between CNS and cardiovascular dysfunction in brain fog. Combined, these approaches offer a roadmap for translating molecular findings into clinical diagnostics and for stratifying long-COVID patients by neurocognitive risk. Further research validating these biomarkers and exploring cardiovascular-focused therapeutic interventions is urgently needed.

The current findings should be interpreted in light of several significant limitations. First, the sample size was modest, which reduced the statistical power and may limit generalizability (Greene *et al*., 2024). Small-cohort RNA□seq studies are especially prone to false negatives and spurious associations; therefore, larger independent cohorts are needed for validation. Ideally, key transcriptomic signals can be confirmed via orthogonal methods (e.g., RT□qPCR or immunoassays) in such cohorts (Grätz *et al*., 2022). Second, we analysed gene expression in peripheral blood, which is only an indirect surrogate for brain physiology. Blood transcriptomes often reflect systemic immune or vascular states, not all of which mirror central nervous system (CNS) processes (Grätz *et al*., 2022; Greene *et al*., 2024). While blood□based biomarkers can track neurologic injury, they are not perfectly concordant with CNS pathology. For example, plasma phosphorylated tau levels have been shown to closely track Alzheimer’s disease pathology (Gonzalez-Ortiz *et al*., 2023). The levels of blood markers of neural injury, such as the neurofilament light chain and GFAP, increase in many neurodegenerative and cerebrovascular conditions (Barro *et al*., 2020) (Grätz *et al*., 2022).

Nevertheless, reliance on blood means that CNS-specific effects may be overlooked. Another limitation is that we used a cross-sectional dataset at a single time point postinfection (∼3–6 months post-COVID-19, according to GEO metadata). We do not know if these gene expression changes are transient or persist (or even worsen) over more extended periods (Chapman *et al*., 2020). Longitudinal studies following the same patients as their brain fog improves or deteriorates would be highly informative in distinguishing cause from effect – do these molecular changes resolve as cognitive function recovers, or do they precede improvement? (Dadsena *et al*., 2025). Furthermore, while we performed pathway analysis, mechanistic insight is still limited; functional studies (in vitro or animal models) are needed to test whether modifying these genes/pathways can influence cognitive outcomes after COVID-19 (McMullen *et al*., 2019).

## 6. Conclusion

Through transcriptomic analysis, this study provides valuable insights into the molecular mechanisms of long COVID-19 brain fog. Patients with brain fog exhibit a distinct gene expression profile characterized by dysregulation of genes involved in neurovascular function, inflammation, and synaptic plasticity. Through transcriptomic analysis, we identified a unique gene expression signature in the brain fog cohort, highlighting genes such as *EDNRB, SLC2A14, SMG1P1, NRCAM*, and the NMDA receptor subunit *GRIN2C*, which were found exclusively in this group. Key findings include a unique molecular signature, neurovascular dysfunction, neuroinflammation, synaptic plasticity alterations, vascular remodelling, and cardiovascular involvement.

This research identifies unique transcriptomic biomarkers and pathways implicating cardiovascular and neurovascular dysregulation in long COVID-related cognitive symptoms. Our findings pave the way for targeted therapeutic strategies addressing these newly emphasized mechanisms, advocating for comprehensive approaches in managing long COVID. Further research is needed to validate these findings in larger cohorts, explore the longitudinal progression of transcriptomic changes, and integrate molecular insights with clinical and neuroimaging data to comprehensively understand long-term COVID-19-related brain fog (Cataldo *et al*., 2024).

## Ethical Statement

This study does not contain any studies with human subjects or animals performed by any of the authors.

## Data availability statement

The current investigation employed information sourced from datasets GEO Accession viewer available from the National Center for Biotechnology Information’s Gene Expression Omnibus (GEO) for the purpose of analysis.

## Acknowledgement

This study was conducted as part of the Research Fellowship Program and received support from OmicsLogic Inc Houstan & OmicsLogic India, New Delhi. The data processing and visualization workflows were executed via the T-Bio Info server.

## Author contributions

Orusa Ashfaque designed the study and performed the formal analysis, investigation, writing and editing of the manuscript. Harsh Kumar was responsible for data extraction and visualization. Mohit Mazumder contributed to study design, manuscript editing, critical review, and providing constructive feedback throughout the manuscript development.

## References

1. Alcendor, D. J. (2020). Dysregulation of endothelin-1: Implications for health disparities in Alzheimer’s disease. Journal of Personalized Medicine, 10(4), 199. 10.3390/jpm10040199

2. Alosco, M. L., Brickman, A. M., Spitznagel, M. B., Garcia, S. L., Narkhede, A., Griffith, E. Y., Raz, N., Cohen, R., Sweet, L. H., Colbert, L. H., Josephson, R., Hughes, J., Rosneck, J., & Gunstad, J. (2013). Cerebral Perfusion is Associated With White Matter Hyperintensities in Older Adults With Heart Failure. Congestive Heart Failure, 19(4). 10.1111/chf.12025

3. Baccari, M., & Bani, D. (2008). Relaxin and nitric oxide signalling. Current Protein and Peptide Science, 9(6), 638–645. 10.2174/138920308786733921

4. Barnden, L., Thapaliya, K., Eaton-Fitch, N., Barth, M., & Marshall-Gradisnik, S. (2023). Altered brain connectivity in Long Covid during cognitive exertion: a pilot study. Frontiers in Neuroscience, 17. 10.3389/fnins.2023.1182607

5. Barro, C., Chitnis, T., & Weiner, H. L. (2020). Blood neurofilament light: a critical review of its application to neurologic disease. Annals of Clinical and Translational Neurology, 7(12), 2508–2523. 10.1002/acn3.51234

6. Bellanti, J. A. (2023). The new frontier: Clinical consequences of long COVID. Journal of Precision Respiratory Medicine, 6(1), 2–9. 10.2500/jprm.2023.6.230001

7. Cataldo, S. A., Micciulli, A., Margulis, L., Cibeyra, M., Defeo, S., Horovitz, S. G., Martino, A., Melano, R., Mena, M., Parisi, F., Santoro, D., Sarmiento, F., & Belzunce, M. A. (2024). Cognitive impact and brain structural changes in long COVID patients: a cross-sectional MRI study two years post infection in a cohort from Argentina. BMC Neurology, 24(1). 10.1186/s12883-024-03959-8

8. Ceban, F., Ling, S., Lui, L. M. W., Lee, Y., Gill, H., Teopiz, K. M., Rodrigues, N. B., Subramaniapillai, M., Di Vincenzo, J. D., Cao, B., Lin, K., Mansur, R. B., Ho, R. C., Rosenblat, J. D., Miskowiak, K. W., Vinberg, M., Maletic, V., & McIntyre, R. S. (2021). Fatigue and cognitive impairment in Post-COVID-19 Syndrome: A systematic review and meta-analysis. Brain Behavior and Immunity, 101, 93–135. 10.1016/j.bbi.2021.12.020

9. Cespuglio, R., Strekalova, T., Spencer, P. S., Román, G. C., Reis, J., Bouteille, B., & Buguet, A. (2021). SARS-CoV-2 infection and sleep disturbances: nitric oxide involvement and therapeutic opportunity. SLEEP, 44(3). 10.1093/sleep/zsab009

10. Chapman, M. A., Arif, M., Emanuelsson, E. B., Reitzner, S. M., Lindholm, M. E., Mardinoglu, A., & Sundberg, C. J. (2020). Skeletal Muscle Transcriptomic Comparison between Long-Term Trained and Untrained Men and Women. Cell Reports, 31(12), 107808. 10.1016/j.celrep.2020.107808

11. Chatys-Bogacka, Ż., Mazurkiewicz, I., Słowik, J., Słowik, A., Drabik, L., & Wnuk, M. (2024). Association between brain fog, cardiac injury, and quality of life at work after hospitalization due to COVID-19. Medycyna Pracy. 10.13075/mp.5893.01428

12. Chen, B. P., & Lane, T. E. (2002). Lack of nitric oxide synthase type 2 (NOS2) results in reduced neuronal apoptosis and mortality following mouse hepatitis virus infection of the central nervous system. Journal of NeuroVirology, 8(1), 58–63. 10.1080/135502802317247820

13. Cheng, Z., Zhang, C., Zhao, F., Piao, J., Cui, R., & Li, B. (2022). RASD2 mediates acute Fasting-Induced Antidepressant-Like effects via dopamine D2 receptor activation in ovariectomized mice. The International Journal of Neuropsychopharmacology, 26(3), 217–229. 10.1093/ijnp/pyac082

14. Chow, B. S. M., Chew, E. G. Y., Zhao, C., Bathgate, R. a. D., Hewitson, T. D., & Samuel, C. S. (2012). Relaxin Signals through a RXFP1-pERK-nNOS-NO-cGMP-Dependent Pathway to Up-Regulate Matrix Metalloproteinases: The Additional Involvement of iNOS. PLoS ONE, 7(8), e42714. 10.1371/journal.pone.0042714

15. Crosier, R., Kafil, T. S., & Paterson, D. I. (2023). Imaging for cardiovascular complications of COVID-19: Cardiac Manifestations in context. Canadian Journal of Cardiology, 39(6), 779–792. 10.1016/j.cjca.2023.01.022

16. Dadsena, R., Walders, J., Costa, A. S., Wetz, S., Romanzetti, S., Lischewski, S. A., Krockauer, C., Heine, J., Schlenker, L., Klabunn, P., Schwichtenberg, K., Hartung, T. J., Franke, C., Balloff, C., Binkofski, F., Schulz, J. B., Finke, C., & Reetz, K. (2025). Two□year impact of COVID□19: Longitudinal MRI brain changes and neuropsychiatric trajectories. Psychiatry and Clinical Neurosciences. 10.1111/pcn.13789

17. Dai, L., Jin, X., & Liu, Z. (2021). Prognostic and clinicopathological significance of GPRC5A in various cancers: A systematic review and meta-analysis. PLoS ONE, 16(3), e0249040. 10.1371/journal.pone.0249040

18. Desai, A. D., Lavelle, M., Boursiquot, B. C., & Wan, E. Y. (2021). Long-term complications of COVID-19. AJP Cell Physiology, 322(1), C1–C11. 10.1152/ajpcell.00375.2021

19. Fernández-Castañeda, A., Lu, P., Geraghty, A. C., Song, E., Lee, M., Wood, J., O’Dea, M. R., Dutton, S., Shamardani, K., Nwangwu, K., Mancusi, R., Yalçın, B., Taylor, K. R., Acosta-Alvarez, L., Malacon, K., Keough, M. B., Ni, L., Woo, P. J., Contreras-Esquivel, D., … Monje, M. (2022). Mild respiratory COVID can cause multi-lineage neural cell and myelin dysregulation. Cell, 185(14), 2452–2468.e16. 10.1016/j.cell.2022.06.008

20. Fontes-Dantas, F. L., Fernandes, G. G., Gutman, E. G., De Lima, E. V., Antonio, L. S., Hammerle, M. B., Mota-Araujo, H. P., Colodeti, L. C., Araújo, S. M., Froz, G. M., Da Silva, T. N., Duarte, L. A., Salvio, A. L., Pires, K. L., Leon, L. A., Vasconcelos, C. C. F., Romão, L., Savio, L. E. B., Silva, J. L., … Figueiredo, C. P. (2023). SARS-CoV-2 Spike protein induces TLR4-mediated long-term cognitive dysfunction recapitulating post-COVID-19 syndrome in mice. Cell Reports, 42(3), 112189. 10.1016/j.celrep.2023.112189

21. Garvin, R. A. (2018). Treatment with relaxin reduces disease symptoms and enhances neuroprotection and remyelination in murine experimental autoimmune encephalomyelitis. Italian Journal of Anatomy and Embryology, 123(1), 64–79. 10.13128/ijae-23012

22. Golzardi, M., Hromić-Jahjefendić, A., Šutković, J., Aydin, O., Ünal-Aydın, P., Bećirević, T., Redwan, E. M., Rubio-Casillas, A., & Uversky, V. N. (2024). The Aftermath of COVID-19: Exploring the Long-Term Effects on Organ Systems. Biomedicines, 12(4), 913. 10.3390/biomedicines12040913

23. González, P. a. I., Valdivieso, Á. G., & Santa-Coloma, T. A. (2023). The G protein-coupled receptor GPRC5A—a phorbol ester and retinoic acid-induced orphan receptor with roles in cancer, inflammation, and immunity. Biochemistry and Cell Biology, 101(6), 465–480. 10.1139/bcb-2022-0352

24. Gonzalez-Ortiz, F., Kac, P. R., Brum, W. S., Zetterberg, H., Blennow, K., & Karikari, T. K. (2023). Plasma phospho-tau in Alzheimer’s disease: towards diagnostic and therapeutic trial applications. Molecular Neurodegeneration, 18(1). 10.1186/s13024-023-00605-8

25. Grätz, C., Bui, M. L. U., Thaqi, G., Kirchner, B., Loewe, R. P., & Pfaffl, M. W. (2022). Obtaining Reliable RT-QPCR Results in Molecular Diagnostics—MIQE Goals and Pitfalls for transcriptional biomarker discovery. Life, 12(3), 386. 10.3390/life12030386

26. Greene, C., Connolly, R., Brennan, D., Laffan, A., O’Keeffe, E., Zaporojan, L., O’Callaghan, J., Thomson, B., Connolly, E., Argue, R., Meaney, J. F. M., Martin-Loeches, I., Long, A., Cheallaigh, C. N., Conlon, N., Doherty, C. P., & Campbell, M. (2024). Blood–brain barrier disruption and sustained systemic inflammation in individuals with long COVID-associated cognitive impairment. Nature Neuroscience, 27(3), 421–432. 10.1038/s41593-024-01576-9

27. Grover, S., Sahoo, S., Mishra, E., Gill, K. S., Mehra, A., Nehra, R., Suman, A., Bhalla, A., & Puri, G. D. (2021). Fatigue, perceived stigma, self-reported cognitive deficits and psychological morbidity in patients recovered from COVID-19 infection. Asian Journal of Psychiatry, 64, 102815. 10.1016/j.ajp.2021.102815

28. Gupta, S. C., Ravikrishnan, A., Liu, J., Mao, Z., Pavuluri, R., Hillman, B. G., Gandhi, P. J., Stairs, D. J., Li, M., Ugale, R. R., Monaghan, D. T., & Dravid, S. M. (2016). The NMDA receptor GluN2C subunit controls cortical excitatory-inhibitory balance, neuronal oscillations and cognitive function. Scientific Reports, 6(1). 10.1038/srep38321

29. Haloot, J., Bhavaraju-Sanka, R., Pillarisetti, J., & Verduzco-Gutierrez, M. (2023). Autonomic dysfunction related to postacute SARS-COV-2 syndrome. Physical Medicine and Rehabilitation Clinics of North America, 34(3), 563–572. 10.1016/j.pmr.2023.04.003

30. Henderson-Smith, A., Chow, D., Meechoovet, B., Aziz, M., Jacobson, S. A., Shill, H. A., Sabbagh, M. N., Caviness, J. N., Adler, C. H., Driver-Dunckley, E. D., Beach, T. G., Yin, H., & Dunckley, T. (2013). SMG1 Identified as a Regulator of Parkinson’s Disease-Associated alpha-Synuclein through siRNA Screening. PLoS ONE, 8(10), e77711. 10.1371/journal.pone.0077711

31. Hon, K. L., Leung, K. K. Y., Leung, A. K., Qian, S. Y., Chan, V. P., Ip, P., & Wong, I. C. (2020). Coronavirus disease 2019 (COVID-19): latest developments in potential treatments. Drugs in Context, 9, 1–14. 10.7573/dic.2020-4-15

32. Hoos, M. D., Vitek, M. P., Ridnour, L. A., Wilson, J., Jansen, M., Everhart, A., Wink, D. A., & Colton, C. A. (2014). The impact of human and mouse differences in NOS2 gene expression on the brain’s redox and immune environment. Molecular Neurodegeneration, 9(1). 10.1186/1750-1326-9-50

33. Iadecola, C., Yaffe, K., Biller, J., Bratzke, L. C., Faraci, F. M., Gorelick, P. B., Gulati, M., Kamel, H., Knopman, D. S., Launer, L. J., Saczynski, J. S., Seshadri, S., & Hazzouri, A. Z. A. (2016). Impact of hypertension on cognitive function: a scientific statement from the American Heart Association. Hypertension, 68(6). 10.1161/hyp.0000000000000053

34. Jiang, X., Xu, X., Wu, M., Guan, Z., Su, X., Chen, S., Wang, H., & Teng, L. (2018). GPRC5A: an emerging biomarker in human cancer. BioMed Research International, 2018, 1–11. 10.1155/2018/1823726

35. Ju, Y., Bai, H., Ren, L., & Zhang, L. (2021). The role of Exosome and the ESCRT pathway on enveloped virus infection. International Journal of Molecular Sciences, 22(16), 9060. 10.3390/ijms22169060

36. Kavanagh, E. (2022). Long Covid brain fog: a neuroinflammation phenomenon? Oxford Open Immunology, 3(1). 10.1093/oxfimm/iqac007

37. Kayalar, O., Cetinkaya, P. D., Eldem, V., Baris, S. A., Kokturk, N., Kuralay, S. C., Rajabi, H., Konyalilar, N., Mortazavi, D., Korkunc, S. K., Erkan, S., Aksoy, G. T., Eyikudamaci, G., Deniz, P. P., Toprak, O. B., Gulhan, P. Y., Sagcan, G., Kose, N., Erdem, A. T., … Bayram, H. (2024). Comparative Transcriptomic Analyses of Peripheral Blood Mononuclear Cells of COVID-19 Patients without Pneumonia and with Severe Pneumonia in the First Year of Follow-Up. Viruses, 16(8), 1211. 10.3390/v16081211

38. Kishor, R. S., & Ramhari, B. M. (2020). Introduction to Covid-19. Research Journal of Science and Technology, 12(4), 338–345. 10.5958/2349-2988.2020.00051.0

39. Koyama, Y. (2021). Endothelin ETB Receptor-Mediated Astrocytic Activation: Pathological roles in brain Disorders. International Journal of Molecular Sciences, 22(9), 4333. 10.3390/ijms22094333

40. Krishnan, K., Lin, Y., Prewitt, K. M., & Potter, D. A. (2022). Multidisciplinary Approach to Brain fog and related Persisting Symptoms Post COVID-19. Journal of Health Service Psychology, 48(1), 31–38. 10.1007/s42843-022-00056-7

41. Lee, C. C. E., Ali, K., Connell, D., Mordi, I. R., George, J., Lang, E. M., & Lang, C. C. (2021). COVID-19-Associated cardiovascular complications. Diseases, 9(3), 47. 10.3390/diseases9030047

42. Lee, J., & Gao, F. (2012). Neuronal functions of ESCRTs. Experimental Neurobiology, 21(1), 9–15. 10.5607/en.2012.21.1.9

43. Li, X., Edén, A., Malwade, S., Cunningham, J. L., Bergquist, J., Weidenfors, J. A., Sellgren, C. M., Engberg, G., Piehl, F., Gisslen, M., Kumlien, E., Virhammar, J., Orhan, F., Rostami, E., Schwieler, L., & Erhardt, S. (2024). Central and peripheral kynurenine pathway metabolites in COVID-19: Implications for neurological and immunological responses. Brain Behavior and Immunity. 10.1016/j.bbi.2024.11.031

44. Lieberman, N. a. P., Peddu, V., Xie, H., Shrestha, L., Huang, M., Mears, M. C., Cajimat, M. N., Bente, D. A., Shi, P., Bovier, F., Roychoudhury, P., Jerome, K. R., Moscona, A., Porotto, M., & Greninger, A. L. (2020). In vivo antiviral host transcriptional response to SARS-CoV-2 by viral load, sex, and age. PLoS Biology, 18(9), e3000849. 10.1371/journal.pbio.3000849

45. López-Hernández, Y., Monárrez-Espino, J., López, D. a. G., Zheng, J., Borrego, J. C., Torres-Calzada, C., Elizalde-Díaz, J. P., Mandal, R., Berjanskii, M., Martínez-Martínez, E., López, J. A., & Wishart, D. S. (2023). The plasma metabolome of long COVID patients two years after infection. Scientific Reports, 13(1). 10.1038/s41598-023-39049-x

46. Low, R. N., Low, R. J., & Akrami, A. (2023). A review of cytokine-based pathophysiology of Long COVID symptoms. Frontiers in Medicine, 10. 10.3389/fmed.2023.1011936

47. Ma, J., Bai, H., Gong, T., Mao, W., Nie, Y., Zhang, X., Da, Y., Wang, X., Qin, H., Zeng, Q., Hu, F., Qi, X., Shi, B., & Zhang, C. (2022). Novel skewed usage of B-cell receptors in COVID-19 patients with various clinical presentations. Immunology Letters, 249, 23–32. 10.1016/j.imlet.2022.08.006

48. Madeira, C., Vargas-Lopes, C., Brandão, C. O., Reis, T., Laks, J., Panizzutti, R., & Ferreira, S. T. (2018). Elevated glutamate and glutamine levels in the cerebrospinal fluid of patients with probable Alzheimer’s disease and depression. Frontiers in Psychiatry, 9. 10.3389/fpsyt.2018.00561

49. Mailian, D. E., & Kolomiets, V. V. (2019). The role of calcium metabolism dysregulation in the pathogenesis of cardiovascular diseases. Russian Journal of Cardiology, 9, 78–85. 10.15829/1560-4071-2019-9-78-85

50. Majeed, A., & Shajar, M. A. (2020). Is hemoglobin the missing link in the pathogenesis of COVID-19? Anaesthesia Pain & Intensive Care, 24(1), 9–12. 10.35975/apic.v24i1.1216

51. Mathew, B., Kumar, R., Harilal, S., M, S., Pappachan, L. K., & PR, R. (2021). Current Perspective of COVID-19 on Neurology: A Mechanistic Insight. Combinatorial Chemistry & High Throughput Screening, 25(5), 763–767. 10.2174/1386207324666210805121828

52. McMullen, P. D., Pendse, S. N., Black, M. B., Mansouri, K., Haider, S., Andersen, M. E., & Clewell, R. A. (2019). Addressing systematic inconsistencies between in vitro and in vivo transcriptomic mode of action signatures. Toxicology in Vitro, 58, 1–12. 10.1016/j.tiv.2019.02.014

53. Michinaga, S., Hishinuma, S., & Koyama, Y. (2023). Roles of astrocytic endothelin ETB receptor in traumatic brain injury. Cells, 12(5), 719. 10.3390/cells12050719

54. Missailidis, D., Ebrahimie, E., Dehcheshmeh, M. M., Allan, C., Sanislav, O., Fisher, P., Gras, S., & Annesley, S. J. (2024). A blood-based mRNA signature distinguishes people with Long COVID from recovered individuals. Frontiers in Immunology, 15. 10.3389/fimmu.2024.1450853

55. Möller, M., Borg, K., Janson, C., Lerm, M., Normark, J., & Niward, K. (2023). Cognitive dysfunction in post□COVID□19 condition: Mechanisms, management, and rehabilitation. Journal of Internal Medicine, 294(5), 563–581. 10.1111/joim.13720

56. Moy, S. S., Nonneman, R. J., Young, N. B., Demyanenko, G. P., & Maness, P. F. (2009). Impaired sociability and cognitive function in Nrcam-null mice. Behavioural Brain Research, 205(1), 123–131. 10.1016/j.bbr.2009.06.021

57. Ohno, M., Sasaki, M., Orba, Y., Sekiya, T., Masum, M. A., Ichii, O., Sawamura, T., Kakino, A., Suzuki, Y., Kida, H., Sawa, H., & Shingai, M. (2021). Abnormal Blood Coagulation and Kidney Damage in Aged Hamsters Infected with Severe Acute Respiratory Syndrome Coronavirus 2. Viruses, 13(11), 2137. 10.3390/v13112137

58. Patel, M. A., Knauer, M. J., Nicholson, M., Daley, M., Van Nynatten, L. R., Martin, C., Patterson, E. K., Cepinskas, G., Seney, S. L., Dobretzberger, V., Miholits, M., Webb, B., & Fraser, D. D. (2022). Elevated vascular transformation blood biomarkers in Long-COVID indicate angiogenesis as a key pathophysiological mechanism. Molecular Medicine, 28(1). 10.1186/s10020-022-00548-8

59. Pulliam, L., Sun, B., Mustapic, M., Chawla, S., & Kapogiannis, D. (2019). Plasma neuronal exosomes serve as biomarkers of cognitive impairment in HIV infection and Alzheimer’s disease. Journal of NeuroVirology, 25(5), 702–709. 10.1007/s13365-018-0695-4

60. Rao, V., & Bhattamisra, S. K. (2020). Recent developments and opportunities in fighting COVID-19. Coronaviruses, 2(7). 10.2174/2666796701999201204120422

61. Rêgo, L. O. S., Braga, L. L. A., Vilas-Boas, G. S., Cardoso, M. S. O., & Duraes, A. R. (2023). Cardiovascular and Neurological Complications of COVID-19: A Narrative review. Journal of Clinical Medicine, 12(8), 2819. 10.3390/jcm12082819

62. Reiken, S., Sittenfeld, L., Dridi, H., Liu, Y., Liu, X., & Marks, A. R. (2022). Alzheimer’s□like signaling in brains of COVID□19 patients. Alzheimer S & Dementia, 18(5), 955–965. 10.1002/alz.12558

63. Reinhold, D., Farztdinov, V., Yan, Y., Meisel, C., Sadlowski, H., Kühn, J., Perschel, F. H., Endres, M., Düzel, E., Vielhaber, S., Guttek, K., Goihl, A., Venø, M., Teegen, B., Stöcker, W., Stubbemann, P., Kurth, F., Sander, L. E., Ralser, M., … Körtvelyessy, P. (2023). The brain reacting to COVID-19: analysis of the cerebrospinal fluid proteome, RNA and inflammation. Journal of Neuroinflammation, 20(1). 10.1186/s12974-023-02711-2

64. Riquelme, P., Amodio, G., Macedo, C., Moreau, A., Obermajer, N., Brochhausen, C., Ahrens, N., Kekarainen, T., Fändrich, F., Cuturi, C., Gregori, S., Metes, D., Schlitt, H. J., Thomson, A. W., Geissler, E. K., & Hutchinson, J. A. (2017). DHRS9 is a stable marker of human regulatory macrophages. Transplantation, 101(11), 2731–2738. 10.1097/tp.0000000000001814

65. Rodriguez-Morales, A. J., Bonilla-Aldana, D. K., Tiwari, R., Sah, R., Rabaan, A. A., & Dhama, K. (2020). COVID-19, an Emerging coronavirus infection: Current scenario and recent developments – An overview. Journal of Pure and Applied Microbiology, 14(1), 05–12. 10.22207/jpam.14.1.02

66. Rubino, E., Italia, M., Giorgio, E., Boschi, S., Dimartino, P., Pippucci, T., Roveta, F., Cambria, C. M., Elia, G., Marcinnò, A., Gallone, S., Rogaeva, E., Antonucci, F., Brusco, A., Gardoni, F., & Rainero, I. (2025). Exome sequencing reveals a rare damaging variant in GRIN2C in familial late-onset Alzheimer’s disease. Alzheimer S Research & Therapy, 17(1). 10.1186/s13195-024-01661-y

67. Saini, G., & Aneja, R. (2021). Cancer as a prospective sequela of long COVID□19. BioEssays, 43(6). 10.1002/bies.202000331

68. Santos-Gómez, A., Miguez-Cabello, F., García-Recio, A., Locubiche-Serra, S., García-Díaz, R., Soto-Insuga, V., Guerrero-López, R., Juliá-Palacios, N., Ciruela, F., García-Cazorla, À., Soto, D., Olivella, M., & Altafaj, X. (2020). Disease-associated GRIN protein truncating variants trigger NMDA receptor loss-of-function. Human Molecular Genetics, 29(24), 3859–3871. 10.1093/hmg/ddaa220

69. Schrank, S., Barrington, N., & Stutzmann, G. E. (2019). Calcium-Handling defects and neurodegenerative disease. Cold Spring Harbor Perspectives in Biology, 12(7), a035212. 10.1101/cshperspect.a035212

70. Serpente, M., Fenoglio, C., Villa, C., Cortini, F., Cantoni, C., Ridolfi, E., Clerici, F., Marcone, A., Benussi, L., Ghidoni, R., Boneschi, F. M., Gallone, S., Cappa, S., Binetti, G., Franceschi, M., Rainero, I., Giordana, M. T., Mariani, C., Bresolin, N., … Galimberti, D. (2011). Role of OLR1 and its regulating HSA-MIR369-3P in Alzheimer’s Disease: Genetics and Expression analysis. Journal of Alzheimer S Disease, 26(4), 787–793. 10.3233/jad-2011-110074

71. Shaghaghi, M. A., Murphy, B., & Eck, P. (2016). The SLC2A14 gene: genomic locus, tissue expression, splice variants, and subcellular localization of the protein. Biochemistry and Cell Biology, 94(4), 331–335. 10.1139/bcb-2015-0089

72. Su, S., Zhao, Y., Zeng, N., Liu, X., Zheng, Y., Sun, J., Zhong, Y., Wu, S., Ni, S., Gong, Y., Zhang, Z., Gao, N., Yuan, K., Yan, W., Shi, L., Ravindran, A. V., Kosten, T., Shi, J., Bao, Y., & Lu, L. (2023). Epidemiology, clinical presentation, pathophysiology, and management of long COVID: an update. Molecular Psychiatry, 28(10), 4056–4069. 10.1038/s41380-023-02171-3

73. Sun, B., Fernandes, N., & Pulliam, L. (2019). Profile of neuronal exosomes in HIV cognitive impairment exposes sex differences. AIDS, 33(11), 1683–1692. 10.1097/qad.0000000000002272

74. Sun, Y., Sun, J., Chen, X., Wang, Y., & Gao, X. (2024). EEG signatures of cognitive decline after mild SARS-CoV-2 infection: an age-dependent study. BMC Medicine, 22(1). 10.1186/s12916-024-03481-1

75. Talkington, G. M., Kolluru, P., Gressett, T. E., Ismael, S., Meenakshi, U., Acquarone, M., Solch-Ottaiano, R. J., White, A., Ouvrier, B., Paré, K., Parker, N., Watters, A., Siddeeque, N., Sullivan, B., Ganguli, N., Calero-Hernandez, V., Hall, G., Longo, M., & Bix, G. J. (2025). Neurological sequelae of long COVID: a comprehensive review of diagnostic imaging, underlying mechanisms, and potential therapeutics. Frontiers in Neurology, 15. 10.3389/fneur.2024.1465787

76. Tugarov, Y., Huet, A., & Dvorshchenko, K. (2023). EXPRESSION OF LRP1 AND OLR1 GENES IN THE BLOOD OF PATIENTSWITH OSTEOARTHRITIS AFTER SARS-COV2 INFECTION. Bulletin of Taras Shevchenko National University of Kyiv Series Biology, 94(3), 35–40. 10.17721/1728.2748.2023.94.35-40

77. Van Der Feltz-Cornelis, C., Turk, F., Sweetman, J., Khunti, K., Gabbay, M., Shepherd, J., Montgomery, H., Strain, W. D., Lip, G. Y., Wootton, D., Watkins, C. L., Cuthbertson, D. J., Williams, N., & Banerjee, A. (2024). Prevalence of mental health conditions and brain fog in people with long COVID: A systematic review and meta-analysis. General Hospital Psychiatry, 88, 10–22. 10.1016/j.genhosppsych.2024.02.009

78. Vishal, N. M. K., U, N. P. W. S., & Kodage, N. P. B. (2024). COVID-19 A comprehensive review of signs, symptoms, diagnosis, and treatment strategies. International Journal of Advanced Research in Science Communication and Technology, 51–68. 10.48175/ijarsct-18110

79. Vitucci, D., Di Giorgio, A., Napolitano, F., Pelosi, B., Blasi, G., Errico, F., Attrotto, M. T., Gelao, B., Fazio, L., Taurisano, P., Di Maio, A., Marsili, V., Pasqualetti, M., Bertolino, A., & Usiello, A. (2015). RASD2 modulates Prefronto-Striatal phenotypes in humans and ‘Schizophrenia-Like behaviors’ in mice. Neuropsychopharmacology, 41(3), 916–927. 10.1038/npp.2015.228

80. Wołoszynowska-Fraser, M. U., Kouchmeshky, A., & McCaffery, P. (2020). Vitamin A and retinoic acid in cognition and cognitive disease. Annual Review of Nutrition, 40(1), 247–272. 10.1146/annurev-nutr-122319-034227

81. Wolters, F. J., Zonneveld, H. I., Hofman, A., Van Der Lugt, A., Koudstaal, P. J., Vernooij, M. W., & Ikram, M. A. (2017). Cerebral perfusion and the risk of dementia. Circulation, 136(8), 719–728. 10.1161/circulationaha.117.027448

82. Ye, Y., Chen, Z., Jiang, S., Jia, F., Li, T., Lu, X., Xue, J., Lian, X., Ma, J., Hao, P., Lu, L., Ye, S., Shen, N., Bao, C., Fu, Q., & Zhang, X. (2022). Single-cell profiling reveals distinct adaptive immune hallmarks in MDA5+ dermatomyositis with therapeutic implications. Nature Communications, 13(1). 10.1038/s41467-022-34145-4

83. Zadeh, A. F., Arnsten, A. F. T., & Wang, M. (2023). Scientific Rationale for the Treatment of Cognitive Deficits from Long COVID. Neurology International, 15(2), 725–742. 10.3390/neurolint15020045

